# The impact of hyperglycemia upon BeWo trophoblast cell metabolic function: A multi-OMICS and functional metabolic analysis

**DOI:** 10.1101/2022.02.15.480604

**Authors:** Zachary JW Easton, Xian Luo, Liang Li, Timothy RH Regnault

**Affiliations:** Department of Physiology and Pharmacology, Western University, Medical Sciences Building Room 216, London, Ontario, Canada, N6A 5C1; The Metabolomics Innovation Centre, University of Alberta, Edmonton, Alberta T6G 2G2, Canada; Department of Chemistry, University of Alberta, Edmonton, Alberta T6G 2G2, Canada; Department of Obstetrics and Gynaecology, London Health Science Centre-Victoria Hospital, B2-401, London, Ontario, Canada, N6H 5W9; Children’s Health Research Institute, 800 Commissioners Road East, London, Ontario, Canada, N6C 2V5; Lawson Health Research Institute, 750 Base Line Rd E, London, Ontario, Canada, N6C 2R5

**Author notes:** Corresponding Author: Dr. Timothy RH Regnault Mailing address: Western University, Dental Science Building, Room 2010, London, Ontario, Canada, N6A 5C1.

## Abstract

Pre-existing and gestationally-developed diabetes mellitus have been linked with impaired metabolic function in placental villous trophoblast cells, that is thought to underlie the development of metabolic diseases early in the lives of the exposed offspring. Previous research using placental cell lines and *ex vivo* trophoblast preparations have highlighted hyperglycemia as an important independent regulator of placental function. However, it is poorly understood if hyperglycemia directly influences aspects of placental metabolic function, including nutrient storage and mitochondrial respiration, that have been found to be impaired in diabetic placentae. Therefore, the current study examined metabolic and mitochondrial function as well as nutrient storage in both undifferentiated cytotrophoblast and differentiated syncytiotrophoblast BeWo cells cultured under hyperglycemia conditions (25 mM glucose) for 72 hours to further characterize the direct impacts of placental hyperglycemic exposure. Hyperglycemic-exposed BeWo trophoblasts displayed increased triglyceride and glycogen nutrient stores. However, specific functional readouts of metabolic enzyme activity and mitochondrial oxidative respiratory activity were not altered. We speculated that increased glycogen content may reduce free glucose available for oxidation and subsequently protect trophoblast cells from mitochondrial damage during acute high glucose exposures. To further characterize the impacts of independent hyperglycemia, the current study subsequently utilized a multi-omics approach and evaluated the transcriptomic and metabolomic signatures of BeWo cytotrophoblasts. Hyperglycemic exposure was associated with an altered transcriptomic profile (e.g. increased expression of *ACSL1* (+1.36 fold-change (FC), *HSD11B2* (+1.35 FC), and *RPS6KA5* (+1.32 FC)), and metabolome profile (e.g. increased levels lactate (+2.72 FC), malonate (+3.74 FC), and riboflavin (+3.68 FC)) in BeWo cytotrophoblasts that further highlighted trophoblast metabolic function is modulated independently by hyperglycemia. Overall, these results demonstrate that hyperglycemia is an important independent regulator of key areas of placental metabolic function and nutrient storage, and these data continue to expand our knowledge on the mechanisms governing the development of placental dysfunction.

## Introduction

The rates of diabetes mellitus (DM) during pregnancy have increased substantially over the past several decades [1]. It is currently estimated that up to 1 in 10 pregnancies worldwide are impacted by maternal DM [2, 3], however these rates may be even higher in certain at risk demographics such as in Indigenous populations [4]. These increases are particularly concerning as maternal DM during pregnancy, regardless of whether it is pre-existing (such as in type 1 DM or type 2 DM) or develops during gestation (gestational DM (GDM)), is associated with a multitude of poor fetal health outcomes [5–8]. Specifically, children exposed to DM during intrauterine development have been found to be at a greater risk of developing non-communicable diseases such as obesity, metabolic syndrome, and impaired insulin sensitivity early in their lives [5,9–12]. Understanding the underlying mechanisms that link maternal DM during pregnancy to the development of non-communicable diseases in offspring is critical to develop appropriate prenatal clinical management practices that help reduce health risks to the next generations.

As the placenta is responsible for nutrient, gas and waste exchange between mother and fetus, specific alterations in the function of this organ may underlie the intrauterine programming of metabolic disorders in DM-exposed offspring. Unsurprisingly, morphological and functional abnormalities have been found to be highly prevalent in placentae of diabetic pregnancies [13, 14]. For example, diabetic placentae are often heavier [15–17], and display increased glycogen and triglyceride content [18–23], that is suggestive of altered nutrient storage and processing by the placenta and subsequently altered nutrient delivery to the developing fetus. This increase in nutrient storage in DM placentae has been thought to modulate trans-placental nutrient transport and fetal growth trajectories [24, 25]. Additionally, the progenitor cytotrophoblasts (CT) and differentiated syncytiotrophoblasts (SCT) cells of the placenta villous trophoblast layer (cells that form the materno-fetal exchange barrier, and are a primary site for placental energy (ATP) production) have been found to have impaired mitochondrial function in response to maternal DM that may further impact placental nutrient handling [26]. In particular, cultured primary CT and SCT cells from GDM pregnancies have been found to have reduced basal and maximal mitochondrial respiratory (oxidative) activity compared to non-diabetic control trophoblasts [27, 28]. Additionally, pre-existing DM has been found to impact the activities of individual placental Electron Transport Chain (ETC) complexes in whole placental lysates, highlighted by reduced complex I, II and III activity in type 1 DM placentae, and reduced complex II and III activity in type 2 DM placentae [29]. Overall, these studies suggest that impaired placental nutrient storage and mitochondrial oxidative function may be implicated in the development of metabolic diseases in DM-exposed offspring.

Previous work with placental cell lines and *ex vivo* placental explant preparations have demonstrated that hyperglycemia (a hallmark symptom of both pre-existing and gestationally-developed DM) is an important regulator of placental metabolic function. For example, explants from uncomplicated pregnancies were found to have altered metabolic processing of lipids when cultured under hyperglycemic (HG) conditions (25 mM glucose) for only 18 hours [22]. Further reports have highlighted transcriptomic and metabolomic markers indicative of altered lipid metabolism, β-oxidation, and glycolysis functions in undifferentiated BeWo CT cells cultured under HG-conditions (25 mM) for 48 hours [30]. Independent hyperglycemia (30 mM glucose for 72h) has also been linked to increased Reactive Oxygen Species (ROS) generation in undifferentiated BeWo CTs [31], which may directly promote the development mitochondrial oxidative damage [32, 33].

These reports have suggested that hyperglycemia independently facilitates the development of abberant placental metabolic function in diabetic pregnancies. However, the direct and independent impacts of hyperglycemia on placental mitochondrial respiratory (oxidative) function are poorly understood. It is important to note that elevated glucose levels (25 mM for 48h) have also been associated with altered mitochondrial activity in BeWo CT cells when assessed by endpoint tetrazolium salt (MTT) assay [34]. However, an interrogation of mitochondrial respiratory activity of HG-exposed trophoblast cells using recently developed real-time functional readouts (such as the Seahorse XF Analyzer), as has been performed with DM-exposed Primary Human Trophoblasts (PHT), [27–29] is warranted. In addition, the direct impacts of hyperglycemia on glycogen and lipid nutrient stores of placental trophoblasts and the underlying mechanisms governing placental nutrient storage in HG-conditions are ill defined. Thus, the first objective of the current study was to characterize the impacts of independent hyperglycemia on placental mitochondrial respiratory activity and nutrient storage by evaluating both undifferentiated BeWo CTs and differentiated BeWo SCTs following a relatively prolonged 72-hour HG (25 mM) exposure.

Recently, the integration of transcriptomics with metabolomics has been identified as a useful method to elucidate cellular mechanisms that underlie pathological placental development in pre-clinical models [30,35,36]. Thus, the second objective of this study was to utilize a multi-omics research approach to thoroughly characterize the underlying mechanisms leading to altered metabolic function in high-glucose exposed BeWo progenitor CT cells. Overall, it was postulated that HG-culture conditions would be associated with increased nutrient storage and impaired mitochondrial respiratory function in BeWo CT and SCT cells, in association with altered transcriptome and metabolome signatures in BeWo CT cells indicative of altered metabolic function.

## Materials and Methods

### Materials

All materials were purchased from Millipore Sigma (Oakville, Canada) unless otherwise specified.

### Cell culture conditions

BeWo (CCL-98) trophoblast cells were purchased from the American Type Culture Collection (ATCC; Cedarlane Labs, Burlington, Canada). Cells were cultured in F12K media (Gibco, ThermoFisher Scientific, Mississauga, Canada) as recommended by the ATCC, and supplemented with 10% Fetal Bovine Serum (Gibco) and 1% Penicillin-Streptomycin (Invitrogen, ThermoFisher Scientific, Mississauga, Canada). All cells were utilized between passages 5-15 and were maintained at 37℃ and 5% CO_2_/95% atmospheric air.

The F12K media contained 7 mM of glucose, a relatively physiological glucose level, and was utilized for low-glucose (LG) controls. F12K media was supplemented to 25 mM glucose for hyperglycemic (HG) culture treatments as previously utilized with BeWo trophoblasts [30, 34]. BeWo trophoblasts cells were plated in LG F12K media at the specifically stated experimental densities and allowed to adhere to cell culture plates overnight before being treated with HG culture media. Cell media was replenished every 24 hours. At T24h and T48h subsets of BeWo trophoblasts were treated with 250 µM 8-Br-cAMP to induce differentiation from cytotrophoblast-like (CT) cells to SCT cells. Cell cultures were collected after 72 hours of high glucose exposure. A schematic of the HG culture protocol is available in Figure 1.

**Figure 1.**
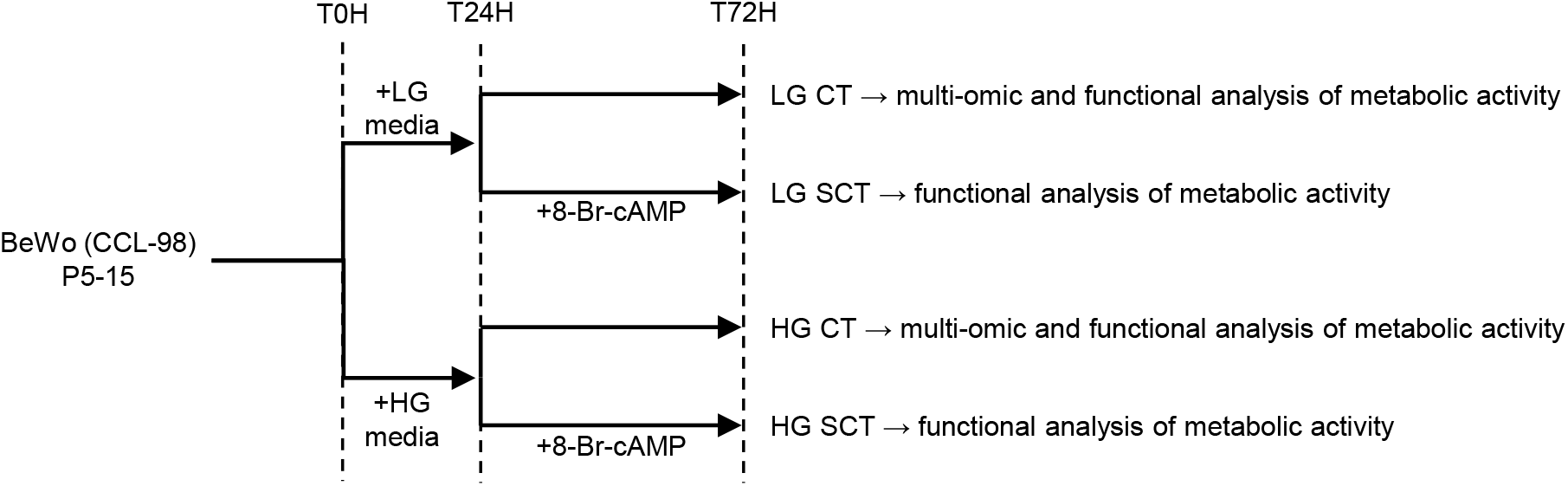
Schematic of 72-hour hyperglycemic cell culture protocol. BeWo trophoblast were plated in F12K media and allowed to adhere to culture plates overnight. At T0H cells were treated with low glucose (LG, 7 mM) or hyperglycemic (HG, 25 mM) supplemented F12K media. At T24H subsets of BeWo trophoblasts were treated with 250 µM 8-Br-cAMP to induce CT-to-SCT differentiation. Cell media was replenished every 24 hours and cells were collected at T72H for analysis of metabolic function.

### Cell Viability of HG-cultured BeWo trophoblasts

BeWo trophoblasts were plated at 7.5×10^3^ cells/well in black walled 96-well cell culture plates and cultured as described. At T72h cell viability of both CT and SCT cultures was assessed via the CellTiter-Fluor cell viability assay (Promega Corporation, Madison WI, USA) as per manufacturer’s instructions.

### Analysis of BeWo cell fusion under HG culture conditions

To determine the potential of HG culture conditions to impact the ability of BeWo trophoblasts to differentiate into SCT cells, cell fusion of 8-Br-cAMP stimulated cells (expressed as percent loss of the tight junction protein zona occludens-1 (ZO-1)) was examined. In brief, BeWo cells were plated at 1.4×10^5^ cells/well in 6-well plates containing coverslips coated with laminin (2 µg/cm^2^) and grown under HG conditions as described. Cellular expression of ZO-1 was then examined via immunofluorescent microscopy as previously detailed [37].

### RT-qPCR Analysis of BeWo syncytialization under HG conditions

The expression of the transcription factor Ovo Like Transcriptional Repressor 1 (OVOL1) as well as human chorionic gonadotropin subunit beta (CGB) was additionally analyzed to ensure cell fusion in 8-Br-cAMP stimulated BeWo cells was associated with increased expression of syncytialization-related genes. In brief, BeWo trophoblasts were plated at 3.5×10^5^ cells/plate in 60mm cell culture plates and cultured as described above. At T72h cells were collected in TRIzol reagent (Invitrogen), and total RNA was extracted as per the manufacturer’s protocol. RNA integrity was assessed via formaldehyde gel electrophoresis, and RNA concentration was quantified by Nanodrop Spectrophotometer 2000 (NanoDrop Technologies, Inc., Wilmington, DE, USA). RNA (2 µg) was then reverse transcribed with the High-Capacity cDNA Reverse Transcription Kit (Applied Biosystems; ThermoFisher Scientific). RT-qPCR was then performed via the CFX384 Real Time system (Bio-Rad, Mississauga, Canada). Relative gene expression of *OVOL1* [38] and *CGB* [39] was then determined using the ΔΔCt method with the geometric mean of *PSMB6* and *ACTB* utilized as a reference gene. Primer sequences and their efficiencies are available in **Supplementary Table 1.**

### Seahorse XF Mito Stress Test quantification of mitochondrial respiratory activity

BeWo cells were plated at 7.5×10^3^ cells/well in Seahorse XF24 V7PS plates and cultured under LG and HG conditions as described above. At T72h mitochondrial activity was assessed using the Seahorse XF Mito Stress Test, as previously optimized for BeWo trophoblast cells [37]. In brief, oxygen consumption rate (OCR) of cell culture media was quantified as a proxy measure of mitochondrial respiratory activity. Subsequent injections of oligomycin (1.5 µg/mL), dinitrophenol (50 µM) and Rotenone and Antimycin A (0.5 µM each) were used to interrogate the basal respiration, maximal respiration, proton leak, spare respiratory capacity and coupling efficiency parameters of mitochondrial respiratory activity. OCR measures were normalized to cellular DNA content using Hoechst dye fluorescence as previously described [37].

### Seahorse XF Glycolysis Stress Test quantification of BeWo trophoblast glycolytic activity

BeWo cells were plated at 7.5×10^3^ cells/well in Seahorse XF24 V7PS plates and cultured under LG and HG conditions as described above. At T72h glycolytic activity was assessed via the Seahorse XF Glycolysis Stress Test, as previously optimized for BeWo trophoblasts [37]. In brief, Extracellular Acidification Rate (ECAR) of cell culture media was assessed to quantify glycolytic activity in BeWo trophoblasts. Injections of glucose (10 mM), oligomycin (1.5 µg/mL) and 2-deoxyglucose (50 mM) were utilized to interrogate the basal glycolytic rate, maximal glycolytic rate, glycolytic reserve, and non-glycolytic acidification parameters of glycolytic function. ECAR data was normalized to cellular DNA content using Hoechst dye fluorescence as previously described [37].

### Immunoblot analysis of protein abundance

BeWo trophoblasts were plated at 9.5×10^5^ cells/plate in 100 mm cell culture dishes and cultured under LG and HG conditions as described. At T72h cells were washed once in ice-cold PBS and subsequently lysed in radioimmunoprecipitation assay (RIPA) buffer supplemented with protease and phosphatase inhibitors as previously described [37]. Protein concentrations were then adjusted to 2 µg/µL in Laemmli loading buffer (62.5 mM Tris-Cl (pH 6.8); 2% SDS 10% glycerol; 0.002% bromophenol blue; 4% β-mercaptoethanol).

The relative abundance of target proteins was then determined via SDS-PAGE gel electrophoresis. In brief, protein lysates were separated with on acrylamide gels (10-15%), and separated proteins were transfer to polyvinylidene fluoride (PVDF) membranes (EMD Millipore, Fisher Scientific). Total lane protein was then determined via Ponceau stain (0.1% Ponceau-S in 5% acetic acid) and utilized to normalize densitometry values. PVDF membranes were blocked in 5% dry-milk protein or 5% Bovine Serum Albumin (BSA, BioShop Canada Inc., Burlington, Canada) and membranes were incubated overnight at 4℃ with respective antibody solutions (**Supplementary Table 2**). Membranes were then washed 3 times with Tris-buffered Saline with 0.1% Tween-20 (TBST) and incubated with respective secondary antibodies for 1 hour at room temperature (**Supplementary Table 2**). Membranes were washed 3 more times in TBST and imaged on a ChemiDoc Imager (Bio-Rad) using Clarity western ECL substrate (Bio-Rad). Protein band abundance and total lane protein (ponceau) were quantified with Image-Lab Software (Bio-Rad).

### Analysis of ETC complex I and II Activity in HG BeWo trophoblasts

BeWo trophoblasts were plated at 3.5×10^5^ cells/plate in 60mm cell culture dishes and cultured under LG and HG conditions as described. At T72h cells were washed once with PBS and detached from cell culture plates by scraping. Cells were then pelleted (400g, 5 mins), snap frozen in liquid nitrogen and stored at −80℃ until analyzed. Cell pellets were subsequently lysed and Complex I activity was assessed as the rate of rotenone-sensitive NADH oxidation, and Complex II activity was assessed as the rate of DCPIP oxidation as previously detailed [37]. ETC complex activity assays were normalized to cell lysate protein content via Bicinchoninic Acid (BCA) assay (Pierce, ThermoFisher Scientific) as per manufacturer’s instructions.

### Analysis of Metabolic Enzyme Activities in HG BeWo trophoblasts

BeWo trophoblasts were plated at 3.5×10^5^ cells/plate in 60mm cell culture dishes and cultured under LG and HG conditions as detailed above. To examine the enzyme activities of Lactate Dehydrogenase (LDH) and Citrate Synthase (CS), cells were collected by scraping and lysed in glycerol lysis buffer (20 mM Na_2_HPO_4_, 0.5 mM EDTA, 0.1% Triton X-100, 0.2% BSA, 50% glycerol) containing protease and phosphatase inhibitors as previously described [37, 40]. LDH activity was assessed as the rate of NADH oxidation, and CS activity was assessed as the rate of Ellman’s reagent consumption as previously detailed [37]. The enzyme activity of LDH and CS were normalized to cell lysate protein content via BCA assay.

Additional BeWo cultures were collected to analyze the activity of the E1 (rate-limiting) subunit of the Pyruvate Dehydrogenase (PDH) complex. PDH-E1 subunit was assessed on freshly collected BeWo cells as the rate of DCPIP oxidation and normalized to protein content via BCA assay as previously detailed [37].

### Nutrient Storage in HG BeWo trophoblasts

BeWo trophoblasts were plated at 3.5×10^5^ cells/plate in 60mm cell culture dishes and cultured under LG and HG conditions. At T72h cells were washed with PBS and collected into fresh PBS (1.5 mL) by scrapping. Cells were then pelleted (400g, 5 minutes) and the PBS was aspirated.

To determine cellular glycogen content, the cell pellets were lysed in 200 µL ddH_2_O and samples were boiled for 10 minutes to inactivate cell enzymes. Samples were stored at −20 until glycogen content was analyzed. Samples were diluted 5-fold and glycogen content was analyzed via the Glycogen Assay Kit (ABCAM, ab65620) as per manufacturer’s protocol. Glycogen content was then normalized to cell lysate protein content via BCA assay.

To determine cellular triglyceride accumulation, the collected cell pellets were snap frozen in liquid nitrogen and stored at −80℃ until analyzed. Cells were then lysed, and cellular triglyceride content was analyzed via the Triglyceride Assay Kit (ABCAM, ab178780) as per kit instructions. Triglyceride content was normalized to cell lysate protein content determined via BCA assay.

### Transcriptomic Analysis of gene expression changes in HG BeWo CT cells

BeWo CT cells were plated at 3.5×10^5^ cells/plate in 60 mm cell culture dishes and grown under LG and HG culture conditions as described. At T72h cells with washed once with PBS and collected in 900 µL TRIzol Reagent and stored at −80℃. Samples were then shipped to the Genome Québec Innovation Centre for transcriptomic analysis via Clariom S mRNA microarray. Automated RNA extraction was completed via QIAcube Connect (Qiagen, Toronto, Canada). RNA content was then quantified via NanoDrop Spectrophotometer 2000 (Nanodrop Technologies, Inc) and RNA integrity was determined by Bioanalyzer 2100 (Agilent Technologies, Waldbronn, Germany). All extracted RNA samples had an RNA Integrity Number (RIN) greater than 9.5. RNA was then processed via the Affymetrix Whole Transcript 2 workflow and analyzed using a Clariom S human mRNA microarray. In brief, sense-stranded cDNA was synthesized from 100 ng total RNA, and subsequently fragmented, and labelled with the GeneChip WT Terminal Labeling Kit (ThermoFisher Scientific) as per manufacturer’s instructions. Labelled DNA was then hybridized to Clariom S human GeneChips (ThermoFisher Scientific) and incubated at 45℃ in the GeneChip Hybridization oven 640 (Affymetrix, ThermoFisher Scientific) for 17 hours at 60 rpm. GeneChips were washed using GeneChip Hybridization Wash and Stain Kit (ThermoFisher Scientific) according to manufacturer’s specifications. Microarray chips were scanned on a GeneChip scanner 3000 (ThermoFisher Scientific).

Microarray data was then analyzed via Transcriptome Analysis Console v4.0 (ThermoFisher Scientific), and raw data was normalized using the Robust Multiple-Array Averaging (RMA) method. HG-treated samples were paired with respective LG-control for each cell collection for analysis. Genes with a ≥ ±1.3 fold-change (FC) vs LG-control and raw-p < 0.05 were determined to be differentially expressed. The list of differentially expressed genes was then imported into WebGestalt for gene ontology (GO) analysis and analysis of functional pathways using the Wikipathways database. Biological processes and functional pathways were determined to be significantly enriched with and a False Discovery Rate (FDR) p < 0.05.

### RT-qPCR Validation of differentially expressed genes identified by mRNA microarray

RT-qPCR was utilized validate differentially expressed genes involved in metabolic pathways that were highlighted by the mRNA microarray. The RNA utilized for the microarray was returned by Genome Québec and 2 µg was reverse transcribed as described above. The CT samples previously utilized to examine expression of syncytialization-related genes were additionally utilized to validate the differentially expressed genes identified in the microarray. RT-qPCR was then performed and analyzed using the ΔΔCt with the geometric mean of *ACTB* and *PSMB6* used as a reference. Primer sequences of validated targets and their efficiencies are available in **Supplementary Table 1.**

### Untargeted Metabolomic Profiling of HG-treated BeWo CT cells

BeWo trophoblasts were plated at a density of 2×10^6^ cells/plate in 150mm cell culture plates and cultured under LG and HG-conditions as described. At T72h cell media was aspirated and the cells were washed three times with cold PBS. Pre-cooled methanol (−20℃) was then added to quench cellular metabolic processes. Cells were then scraped and collected into microcentrifuge tubes, and the methanol was evaporated with a gentle flow of nitrogen gas. Samples were then frozen at −80℃, and subsequently were lyophilized to remove any residual moisture. The samples were then sent to The Metabolomics Innovation Centre (Edmonton, Canada) for subsequent metabolomics analysis via a High-Performance Chemical Isotope Labelling (HP CIL) liquid chromatography mass spectrometry (LC-MS) approach [41, 42].

Samples were reconstituted in 50% methanol and freeze-thaw cycles were utilized to lyse the cells. The lysed samples were centrifuged at 16000 g at 4 ℃ and the supernatants were transferred to new vials. The supernatants were dried down and re-dissolved in 41 µL of water. The total concentrations of metabolite were then determined via the NovaMT Sample Normalization kit (Edmonton, Alberta). Water was added to adjust all the concentrations of samples to 2 mM. The samples were split into five aliquots for respective labeling methods. Each of the individual samples was labeled by ^12^C-DnsCl, base activated ^12^C-DnsCl, ^12^C-DmPA Br, and ^12^C-DnsHz, for amine-/phenol-, hydroxyl-, carboxyl, and carbonyl-metabolomic profiling, respectively [43]. A pooled sample was generated by mixing of each individual sample and labeled by ^13^C-reagent, accordingly. After mixing the each of ^12^C-labeled individual sample with ^13^C-labeled pool by equal volume, the mixtures were injected onto LC-MS for analysis. The LC-MS system was the Agilent 1290 LC (Agilent Technologies) linked to the Bruker Impact II QTOF Mass Spectrometer (Bruker Corporation, Billerica, US). LC-MS data was then exported to .csv files with Bruker DataAnalysis 4.4 (Bruker Corporation), the exported data were then uploaded to IsoMS pro v1.2.7 for data quality check and data processing.

Metabolite peak pairs were then identified using a three-tier approach [43]. In tier 1, peak pairs were identified by searching against a labelled metabolite library (CIL library) based on accurate mass and retention time. In tier 2, the remaining peak pairs were matched by searching against a linked identity library (LI library), containing predicted retention time and accurate mass information. In tier 3, the rest of peak pairs were matched by searching against MyCompoundID (MCID) library, containing accurate mass information of metabolites and their predicted products Metabolites with ± 1.5 fold-change and False Discovery Rate (FDR)-adjusted p<0.05 vs LG CT were determined to be differentially abundant in the HG-cultured BeWo CT cells. Subsequent pathway analysis (Homo sapiens KEGG library) was performed using MetaboAnalyst v5.0 to elucidate the biological impacts of the differentially abundant metabolites identified in tiers 1 and 2. Pathways with an FDR p<0.05 were determined to be significantly enriched.

### Statistical Analysis

Data collected as a percentage (percent loss of ZO-1 staining data, spare respiratory capacity, coupling efficiency, and glycolytic reserve) were log-transformed and analyzed via Two-Way ANOVA (2WA) and Bonferroni’s Multiple Comparisons post-hoc test. A Randomized Block Design 2WA and Sidak’s Multiple Comparisons Test was utilized to analyze relative transcript abundances; relative protein abundances; metabolic activity parameters; and nutrient storage data, using raw data with data from each experimental replicate blocked together, as previously described [44]. These data were then expressed as percent of LG CT control for visualization in figures. Statistical analysis was performed with GraphPad Prism 8 Software (GraphPad Software, San Diego, CA, USA). A paired T-test was utilized to analyze gene FC data between HG and LG BeWo CT cells in the RT-qPCR validation of the microarray.

## Results

### Characterization of BeWo viability and differentiation under high glucose culture conditions

HG culture conditions were associated with a mean 7% and 10% reduction in cell viability in BeWo CT and SCT cells respectively (Fig. 2; p<0.01; n=6/group). It is important to note that these viability data were consistent with previously reported values in BeWo trophoblasts and PHTs cultured under hyperglycemia (25 mM glucose) for 48h [30, 45]. This highlights that a 72H HG-culture protocol does not impact the stability of cultured BeWo trophoblasts cells. Additionally, BeWo SCT cells displayed lower viability relative to BeWo CT cells consistent with trophoblast cells becoming less proliferative while undergoing syncytialization (Fig. 2; p<0.001).

**Figure 2.**
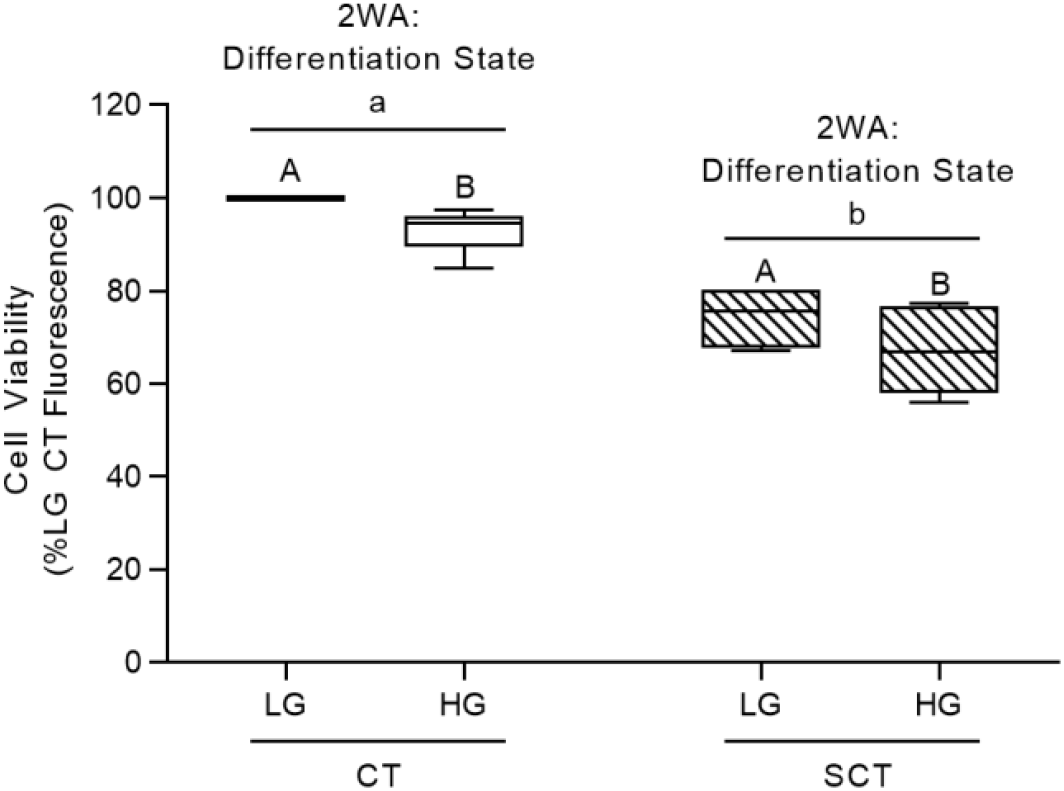
Viability of BeWo trophoblasts cultured under hyperglycemic (HG) conditions for 72h. BeWo trophoblasts were cultured for 72H under HG culture conditions as described, and cell viability was assessed via the CellTiter-Flour cell viability assay. Data is presented as percent of LG CT cell viability (n=6/group). Raw cell viability fluorescence data was analyzed via Two-Way Randomized block design ANOVA (2WA). Different lower-case letters denote differentiation state-dependent differences in viability, and different upper-case letter denote differences in viability within each differentiation state (p<0.05).

BeWo SCT cultures displayed a greater loss of ZO-1 protein expression (and thus increased cell fusion) compared to BeWo CT cultures (Fig 3A,B; p<0.0001, n=4/group). However, HG-culture conditions had no impact on ZO-1 protein expression in BeWo trophoblast cells (Fig 3A,B).

**Figure 3.**
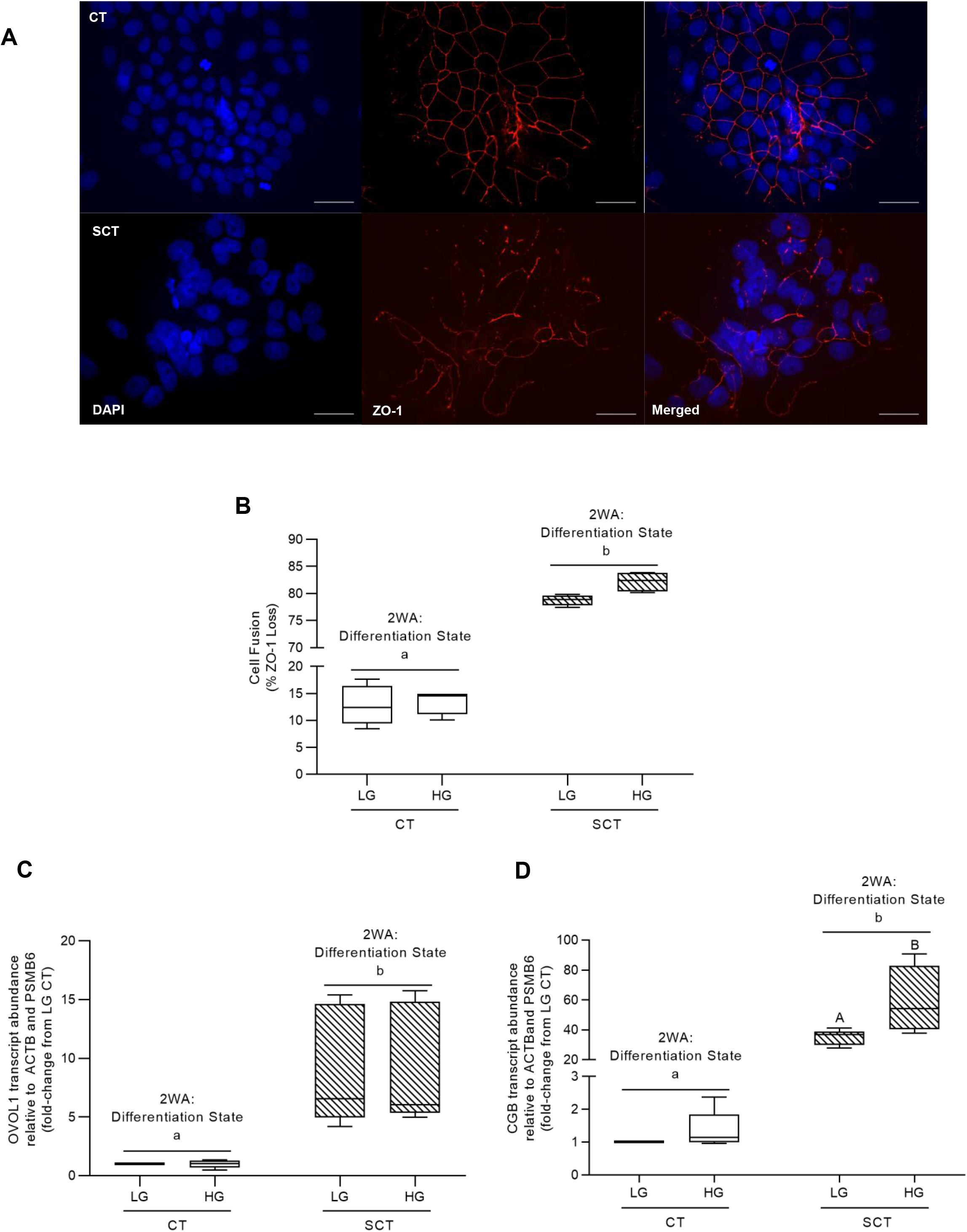
Hyperglycemic (HG) culture conditions do not impact BeWo SCT cell fusion but are associated with increased CGB transcript abundance at 72h. **(A)** Representative DAPI (blue); Zona occludens-1 (ZO1; red) and merged immunofluorescent images (scale bar = 50 µm) of BeWo CT and SCT cells. **(B)** Percent fusion of HG-cultured BeWo trophoblasts. Percent fusion data was log-transformed and analyzed via Two-Way Anova (2WA); data is expressed as the percentage of cells lacking ZO-1 expression (n=4/group). BeWo cell syncytialization was additionally analyzed via quantifying mRNA transcript abundance of the syncytialization markers **(C)** OVOL1 and **(D)** CGB. Transcript abundance data was analyzed via Randomized Block 2WA and Sidak’s multiple comparisons test; data is present as transcript FC versus LG CT cultures (n=5/group). Different lower-case letters denote statistical differences between differentiation states, and different upper-case letter denote differences between LG and HG treatments within each respective differentiation state (p<0.05).

BeWo SCT cultures additionally displayed increased relative transcript abundance of the syncytialization-associated transcription factor OVOL1 (Fig. 3C; p<0.01, n=5/group) as well as the syncytialization-associated hormone CGB (Fig. 3D; p<0.01, n=5/group). HG culture conditions in BeWo SCT cells was additionally associated with an increased transcript abundance of CGB compared to LG BeWo SCT cells (Fig. 3D; p<0.05; 2WA: Interaction p<0.05).

### BeWo mitochondrial respiratory and glycolytic activity

High glucose culture conditions did not impact any of the parameters of mitochondrial respiratory function as assessed by the Seahorse XF Mito Stress Test (Table 1; n=5/group). However, BeWo syncytialization was associated with reduced Spare Respiratory Capacity (Table 1; p<0.0001) and reduced Coupling Efficiency (Table 1; p<0.01) independent of culture glucose level.

**Table 1:** HG culture conditions do not impact BeWo trophoblast glycolytic activity.

| Differentiation State | Treatment | Basal glycolytic Rate (ECAR/μg DNA) | Max Glycolytic Rate (ECAR/μg DNA) | Reserve Glycolytic Capacity (% basal glycolysis) | Non-Glycolytic Acidification (ECAR/μg DNA) |
| --- | --- | --- | --- | --- | --- |
| CT | LG | 75.71 ± 15.38 | 101.4 ± 20.92 | 133.8 ± 1.14 | 19.26 ± 4.43 |
| | HG | 67.54 ± 7.30 | 88.23 ± 11.82 | 129.5 ± 4.08 | 18.38 ± 3.10 |
| SCT | LG | 69.21 ± 12.67 | 84.45 ± 14.55 | 123.5 ± 3.70 | 20.50 ± 3.39 |
| | HG | 77.38 ± 6.78 | 88.01 ± 8.17 | 113.6 ± 1.45 | 23.30 ± 2.45 |
| *Differentiation state difference |  | NS | NS | * | NS |

Data are mean ± SEM (n=5/group). \* denotes a differentiation state dependent difference in glycolytic activity between BeWo CT and SCT cells

The parameters of glycolytic function as measured with the Seahorse XF Glycolysis Stress Test were not impacted by BeWo syncytialization, or by HG culture conditions (Table 2).

**Table 2:** Syncytialization but not HG culture conditions impacts BeWo trophoblast mitochondrial respiratory activity.

| Differentiation State | Treatment | Basal Respiration (OCR/μg DNA) | Maximal Respiration (OCR/μg DNA) | Proton Leak (OCR/μg DNA) | Spare Respiratory Capacity (% Basal OCR) | Coupling Efficiency (% Basal OCR) |
| --- | --- | --- | --- | --- | --- | --- |
| CT | LG | 199.70 ± 39.08 | 249.10 ± 43.78 | 55.48 ± 8.45 | 129.00 ± 5.30 | 70.38 ± 3.34 |
| | HG | 173.40 ± 24.69 | 221.30 ± 36.78 | 46.72 ± 3.76 | 125.50 ± 6.79 | 70.58 ± 4.59 |
| SCT | LG | 173.00 ± 7.96 | 154.40 ± 11.97 | 78.20 ± 7.38 | 89.02 ± 5.68 | 55.04 ± 2.99 |
| | HG | 177.5 ± 19.07 | 168.30 ± 18.27 | 75.88 ± 12.12 | 95.76 ± 5.83 | 57.30 ± 3.80 |
| *Differentiation state difference |  | NS | NS | NS | * | * |

Data are mean ± SEM (n=5/group). \* denotes a differentiation state dependent difference in mitochondrial respiratory activity between BeWo CT and SCT cells

### The impact of high glucose and syncytialization upon BeWo ETC complex protein abundance and activity

High glucose levels did not impact protein abundance of ETC complex subunits in BeWo trophoblasts (Fig. 4A-E; n=4-5/group). Likewise, HG-culture conditions did not affect ETC complex I or II activity in BeWo trophoblasts (Table 3).

**Fig. 4.**
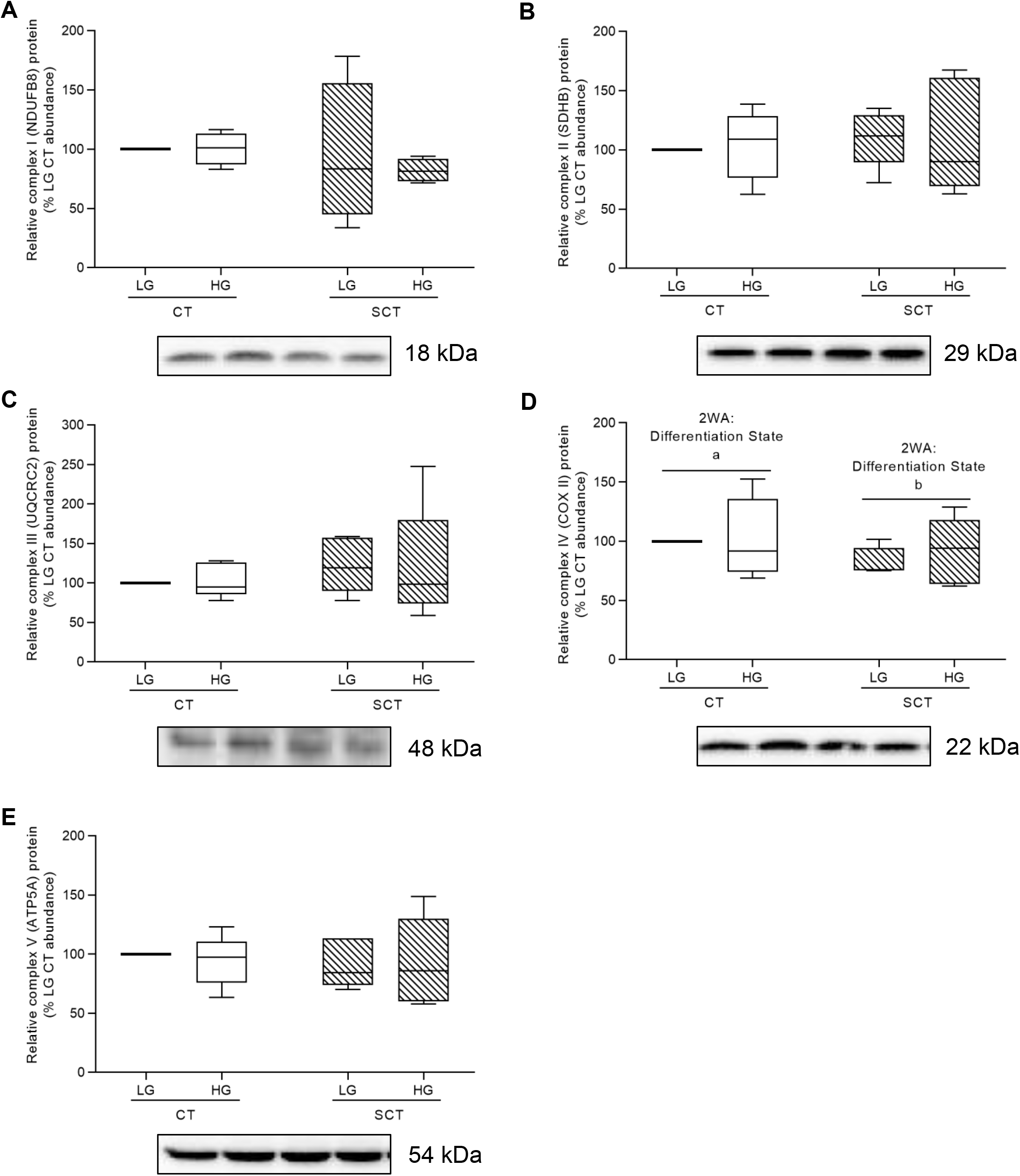
Hyperglycemic (HG) culture conditions do not impact protein expression of Electron Transport Chain (ETC) complexes in BeWo trophoblasts. Relative protein abundance of **(A)** Complex I (NDUFB8 subunit) **(B)** Complex II (SDHB subunit) **(C)** Complex III (UQCRC2 subunit) **(D)** Complex IV (COX II subunit), and (E) Complex V (ATP5A subunit) of the ETC in HG-cultured BeWo trophoblasts. Different lower-case letters denote differentiation state-dependent differences in ETC complex protein abundance (n=4-5/group; Two-Way Randomized Block ANOVA (2WA)). ETC complex protein band density was normalized to total lane protein (ponceau) for statistical analysis and the data is presented as percent of LG CT protein abundance for visualization. Full length representative images of ETC complex bands and ponceau staining of total lane protein are available in **Supplementary Figure 2.**

**Table 3:** Syncytialization but not HG-culture media impacts BeWo metabolic enzyme activity and individual ETC complex activity

| Differentiation State | Treatment | Citrate Synthase | ETC Complex I | ETC Complex II (U/mg protein) | Lactate Dehydrogenase | PDH-E1 subunit |
| --- | --- | --- | --- | --- | --- | --- |
| CT | LG | 0.0126 ± 0.0034 | 0.0354 ± 0.0040 | 0.0470 ± 0.0036 | 2.821 ± 0.277 | 0.0046 ± 0.0006 |
| | HG | 0.0134 ± 0.0033 | 0.0413 ± 0.0052 | 0.0448 ± 0.0039 | 3.095 ± 0.391 | 0.0041 ± 0.0005 |
| SCT | LG | 0.0080 ± 0.0020 | 0.0722 ± 0.0208 | 0.0321 ± 0.0035 | 3.589 ± 0.312 | 0.0090 ± 0.0012 |
| | HG | 0.0070 ± 0.0015 | 0.0709 ± 0.0217 | 0.0330 ± 0.0039 | 3.504 ± 0.359 | 0.0098 ± 0.0014 |
| *Differentiation state difference |  | NS | NS | * | NS | * |

Data are mean ± SEM (n=5-6/group). \* denotes a differentiation state dependent difference in enzyme activity between BeWo CT and SCT cells.

However, BeWo syncytialization was associated with reduced ETC complex IV Cytochrome c oxidase subunit II (COXII) relative protein abundance in both LG and HG cultures (Fig. 4D; p<0.05). Additionally, BeWo SCT cells displayed reduced activity of ETC complex II compared to CT cells independent of culture glucose level (Table 3; p<0.001).

### HG-cultured BeWo trophoblast mitochondrial fission and fusion dynamics

Regardless of glucose level BeWo SCT cells displayed increased relative protein abundance of the mitochondrial fission marker DRP1 in conjunction with decreased relative protein levels of the mitochondrial fusion marker OPA1 compared to undifferentiated BeWo CT cells (Fig. 5A,C; p<0.05, n=5/group).

**Figure 5.**
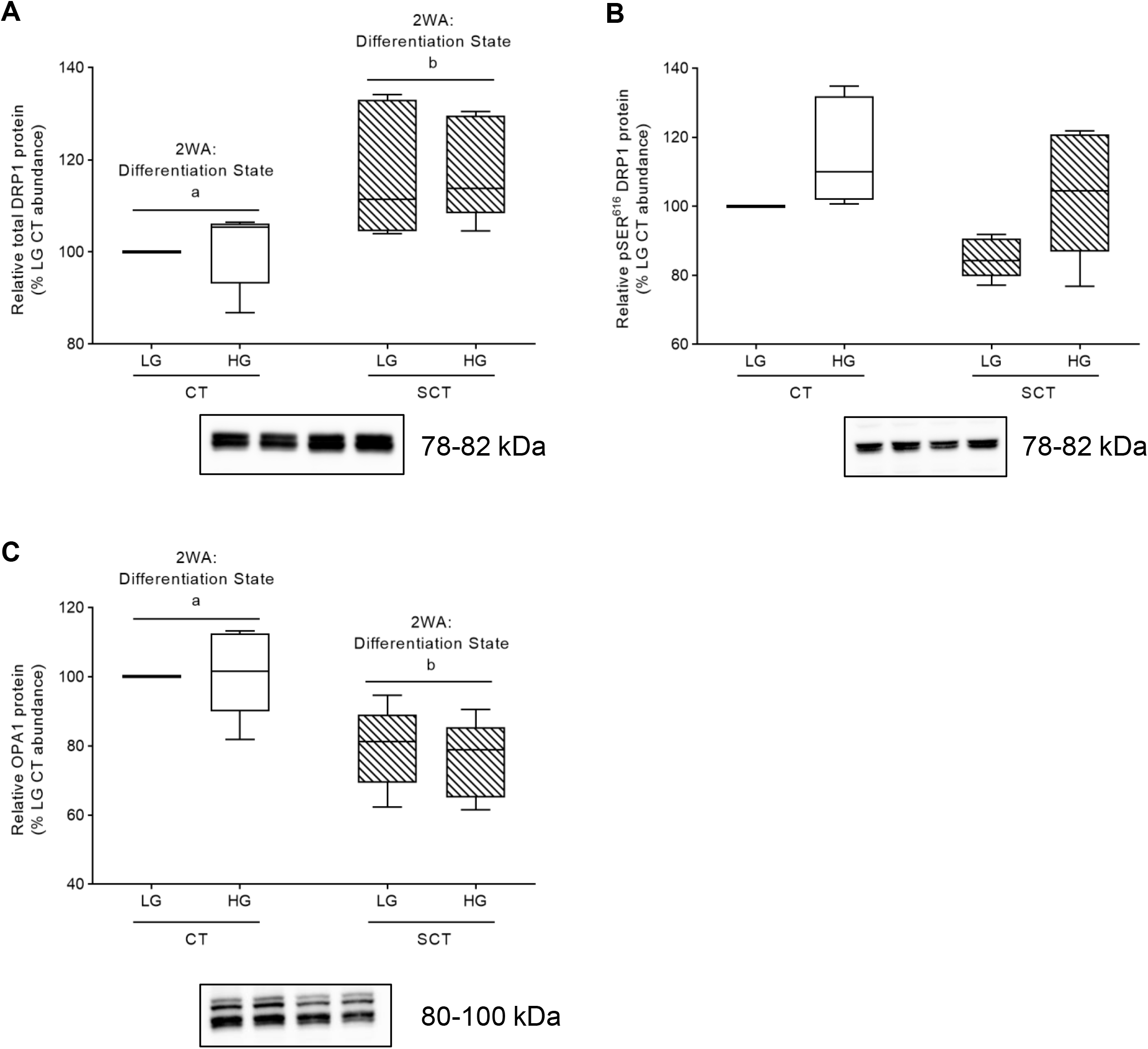
Syncytialization impacts mitochondrial dynamics in BeWo trophoblasts. Relative protein abundance of **(A)** total DRP1 **(B)** pSER616 DRP1 **(C)** OPA1. Different lower-case letters denote differentiation state-dependent differences in relative protein abundance (n=5/group; Two-Way Randomized Block ANOVA (2WA)). Protein band density was normalized to total lane protein (ponceau) for statistical analysis and the data is presented as percent of LG CT abundance for visualization. Uncropped representative images of protein bands and ponceau staining of total lane protein are available in **Supplementary Figures 3-5.**

HG culture conditions additionally did not impact total protein abundance of OPA1 and DRP1 in BeWo trophoblasts (Fig. 5A,C). However, there was a trend towards increased pSER^616^ phosphorylation of DRP1 in HG-cultured BeWo trophoblasts (mean 19% increase; Fig. 5B; p=0.0632, n=5/group) as well as reduced pSER^616^ DRP1 phosphorylation in syncytialized BeWo trophoblasts (Fig. 5B; p=0.0924).

### Metabolic enzyme activity and HG-cultured BeWo trophoblasts

BeWo SCT cells displayed increased activity of the PDH-E1 subunit compared to BeWo CT cells regardless of glucose level (Table 3; p<0.01). Additionally, high glucose levels did not impact the activities of the PDH-E1 subunit, citrate synthase (CS) or lactate dehydrogenase (LDH) in BeWo CT and SCT cells (Table 3). Syncytialization additionally did not affect the enzyme activities of CS or LDH in BeWo trophoblasts (Table 3).

### HG culture conditions impact glycogen storage in BeWo trophoblasts

HG culture conditions in both BeWo CT and SCT cells resulted in increased cellular glycogen content compared to respective differentiation state LG cultures (Fig. 6A; p<0.0001; n=4/group). Furthermore, the glycogen content in BeWo CT cultures was found to be greater than that of the SCT cultures (Fig. 6A; p<0.0001). In addition, BeWo SCT cells displayed increased GLUT1 protein abundance compared to BeWo CT cells (Fig. 6B; p<0.05), however, no glucose-dependent impacts to GLUT1 protein abundance were observed.

**Figure 6.**
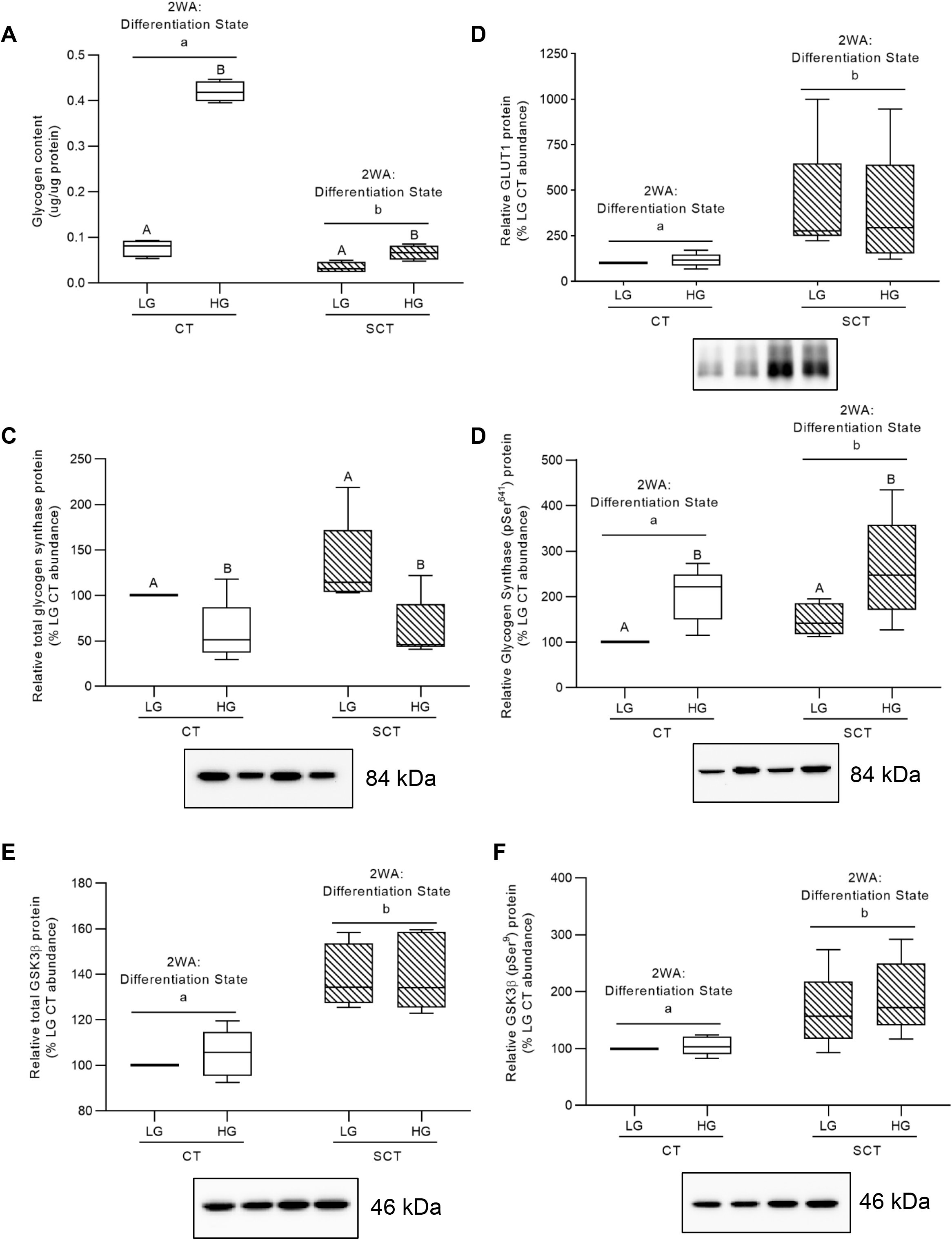
Hyperglycemic (HG) culture conditions impact glycogen storage and regulation in BeWo trophoblasts. **(A)** Glycogen content and relative protein abundance of **(B)** GLUT1 **(C)** glycogen synthase; **(D)** pSer^641^ glycogen synthase; **(E)** GSK3β; **(F)** pSer^9^ GSK3β in HG-treated BeWo trophoblasts at T72H. Different lower-case letters denote differentiation state-dependent differences in protein abundance, and different upper-case letters denote glucose-level dependent differences within a differentiation state (Randomized-Block Two-Way ANOVA (2WA), and Sidak’s multiple comparisons test, p<0.05). Glycogen content data are presented as protein normalized glycogen abundance (µg glycogen per µg protein; n=4/group). Protein band density was normalized to total lane protein (ponceau) for statistical analysis and the data is presented as percent of LG CT abundance for visualization (n=5/group). Uncropped representative images of protein bands and ponceau staining of total lane protein are available in **Supplementary Figures 6-8.**

However, HG culture conditions were associated with reduced relative glycogen synthase protein abundance in both BeWo CT and SCT cells (Fig. 6C; p<0.05, n=5/group). HG cultured BeWo CT and SCT cells subsequently displayed increased relative abundance of phosphorylated (pSer^641^) glycogen synthase (Fig. 6D; p<0.01, n=5/group), and differentiated BeWo SCT cells displayed increased phosphorylated (pSer^641^) glycogen synthase levels relative to undifferentiated CT cultures (Fig. 6D; p<0.01, n=5/group).

Furthermore, differentiated BeWo SCT cells were found to have increased protein levels of both GSK3β and phosphorylated (pSer^9^) GSK3β relative to BeWo CT cultures (Fig. 6E-F; n=5/group). However, HG culture conditions were not associated with alterations in the relative protein abundance of GSK3β and phosphorylated (pSer^9^) GSK3β (Fig. 6E-F; n=5/group).

### HG culture conditions increases TG abundance in BeWo CT cells

HG culture conditions were associated with increased triglyceride accumulation in BeWo CT cells, but not in BeWo SCT cells (Fig. 7A; p<0.01, n=4/group). Furthermore, BeWo syncytialization and HG culture conditions did not impact the relative abundance of ACSL1 protein, although there was a trend towards increased expression in both differentiated BeWo SCT cells and in HG-treated BeWo trophoblasts (Fig. 7B; 2WA: glucose level p=0.0894; 2WA: differentiation state p=0.0695, n=5/group). Finally, syncytialization and HG culture conditions did not impact the relative abundance of fatty acid synthase (FASN) in BeWo trophoblasts (Fig. 7C).

**Figure 7.**
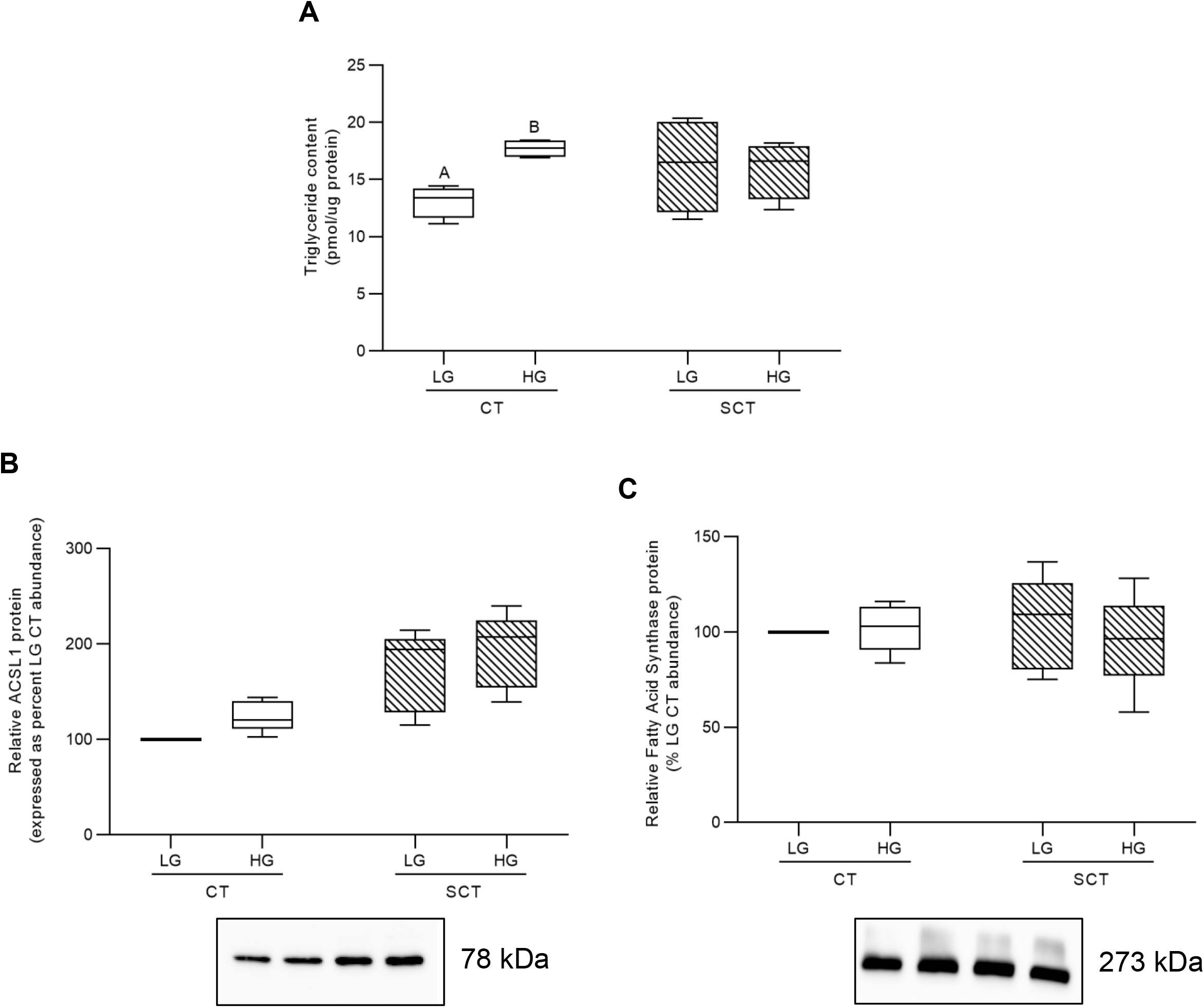
Hyperglycemic (HG) culture conditions impact triglyceride content in BeWo CT cells. **(A)** Triglyceride content and relative protein abundance of **(B)** ACSL1, and **(C)** FASN in HG-treated BeWo trophoblasts at T72H relative abundance. Different upper-case letters denote differences between LG and HG treatments within each respective differentiation state (Two-Way Randomized Block ANOVA; Sidak’s multiple comparisons test, p<0.05). TG content data is presented as protein normalized TG abundance (pmol TG per µg protein; n=4/group). Protein band density was normalized to total lane protein (ponceau) for statistical analysis and the data is presented as percent of LG CT abundance for visualization (n=5/group). Uncropped representative images of protein bands and ponceau staining of total lane protein are available in **Supplementary Figures 3 and 9.**

### Transcriptomic profiling of HG-cultured BeWo CT cells

HG-cultured BeWo CT cells displayed 197 differentially expressed genes (75 upregulated, 122 down-regulated) compared to LG BeWo CT cells (**Supplementary Table 3,** ≥ ±1.3-FC vs LG CT). The volcano plot (Fig. 8) and 2D hierarchical clustering heatmap (Fig. 9) were constructed to visualize the degree of gene expression differences between LG and HG cultured BeWo CT cells.

**Figure 8.**
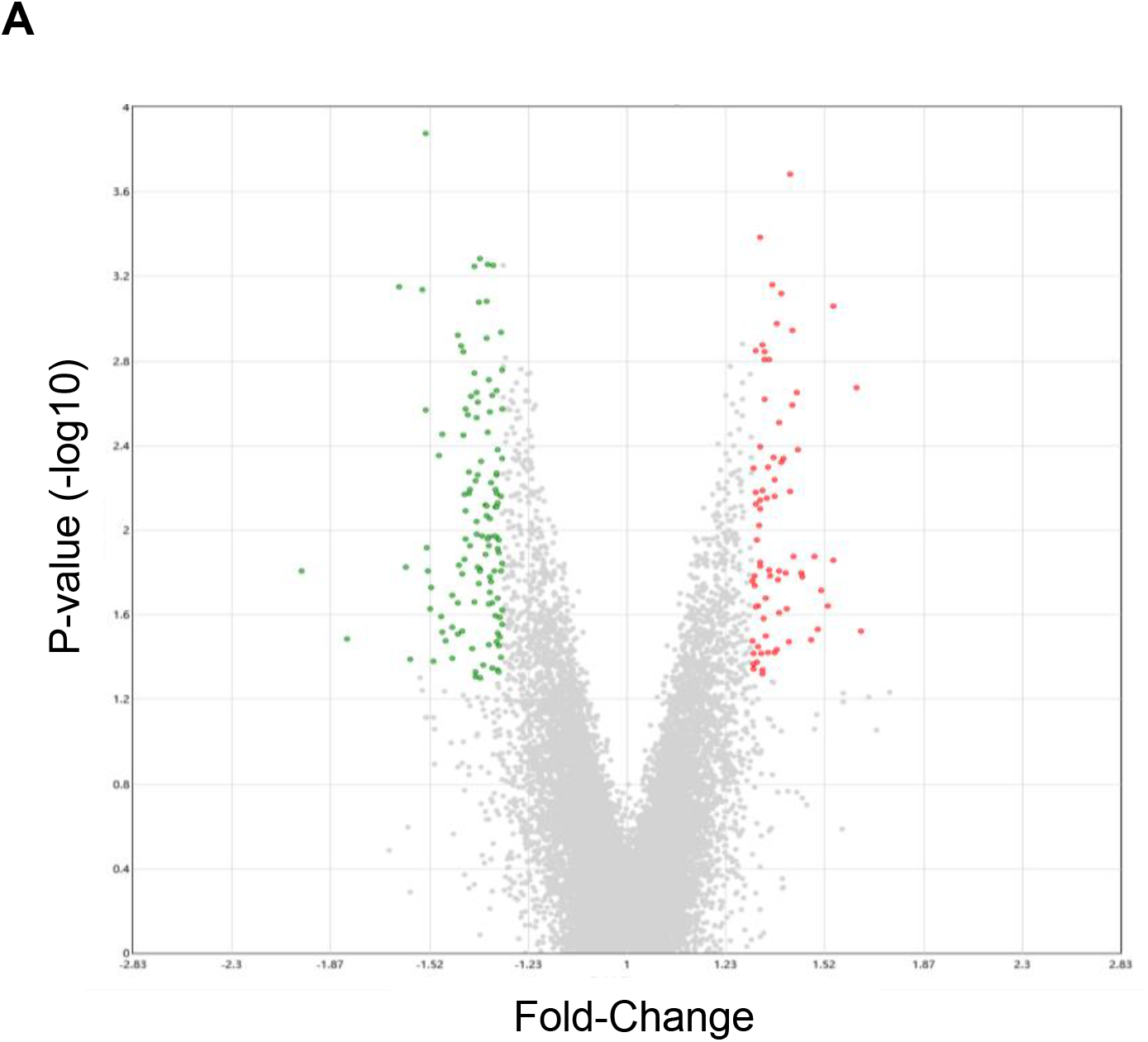
Volcano plot visualized differentially expressed genes between low-glucose and hyperglycemic (HG) cultured BeWo CT cells. The volcano plot was generated to visualized differentially expressed genes in HG-cultured BeWo CT cells (≥ ±1.3-fold change, p<0.05, n=5/group). The x-axis indicated fold-changes vs LG BeWo CTs, and the y-axis indicates the p-value (−log10). The green dots represent statistically significant down-regulated genes, and the red dots represent statistically significant upregulated genes in HG-cultured BeWo CT cells.

**Figure 9.**
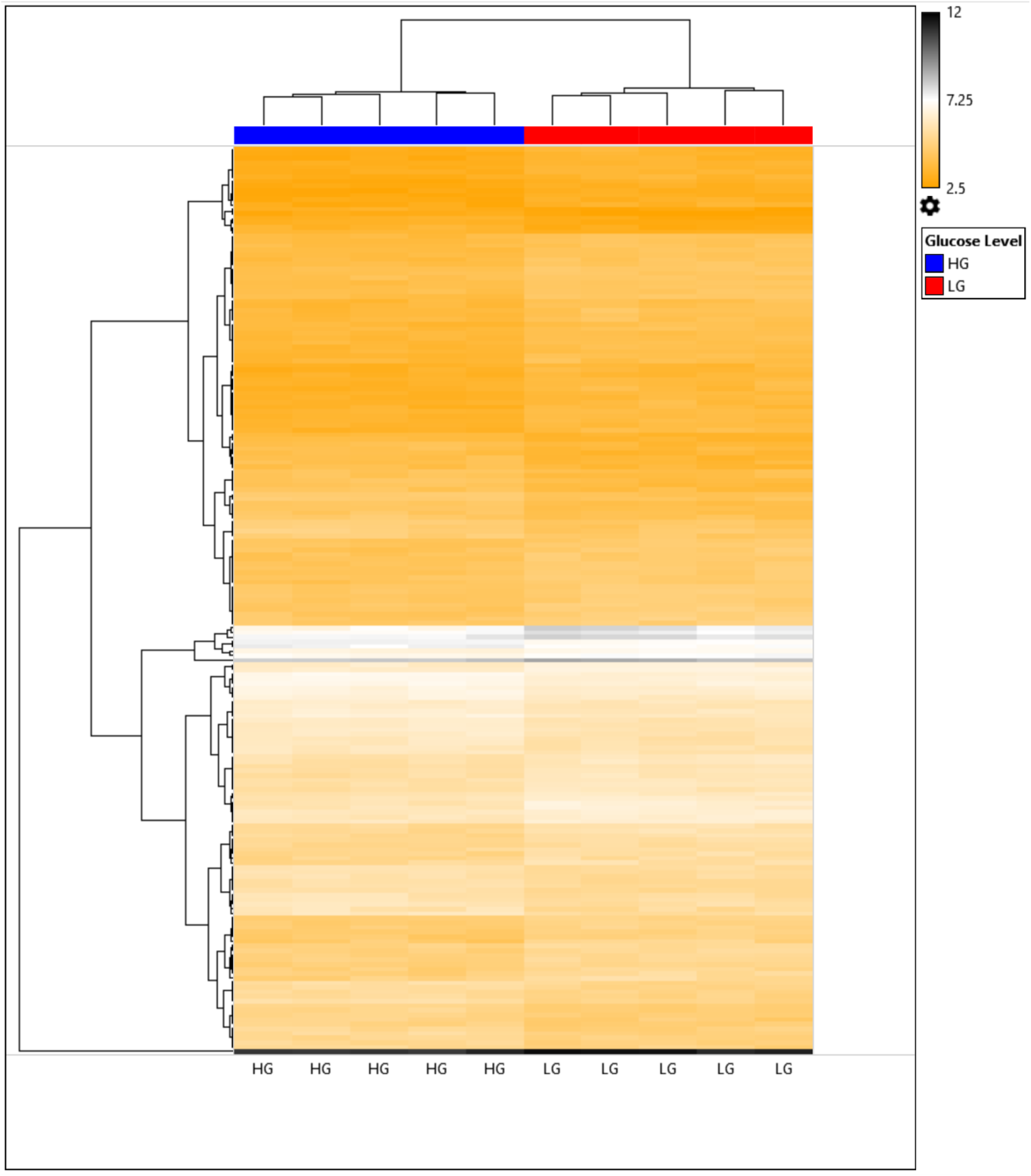
Heat map visualization of differentially expressed genes in hyperglycemic (HG) cultured BeWo CT cells. Each column of the heat map represents an individual sample, with blue columns representing HG-cultured BeWo CT cells and the red columns representing low-glucose (LG) cultured BeWo CT cells. Each row represents an individual differentially expressed gene. Gene expression intensity is color coded with orange representing low-expression genes and black representing high-expression genes.

At the FC cut-offs selected, the differentially expressed genes were not associated with any significant enrichment in gene ontology (GO) biological processes (Fig. 10A). Functional pathway analysis with WikiPathways revealed that the Overview of Nanoparticle Effects Pathway was significantly enriched in the HG-cultured BeWo CT cells (Fig. 10B; FDR < 0.05). The differentially expressed genes involved in this pathway were: *CCND3* (−1.41 FC), *IL6* (−1.39 FC), *PTGS1* (−1.31 FC), and *PTK2* (−1.40 FC).

**Figure 10.**
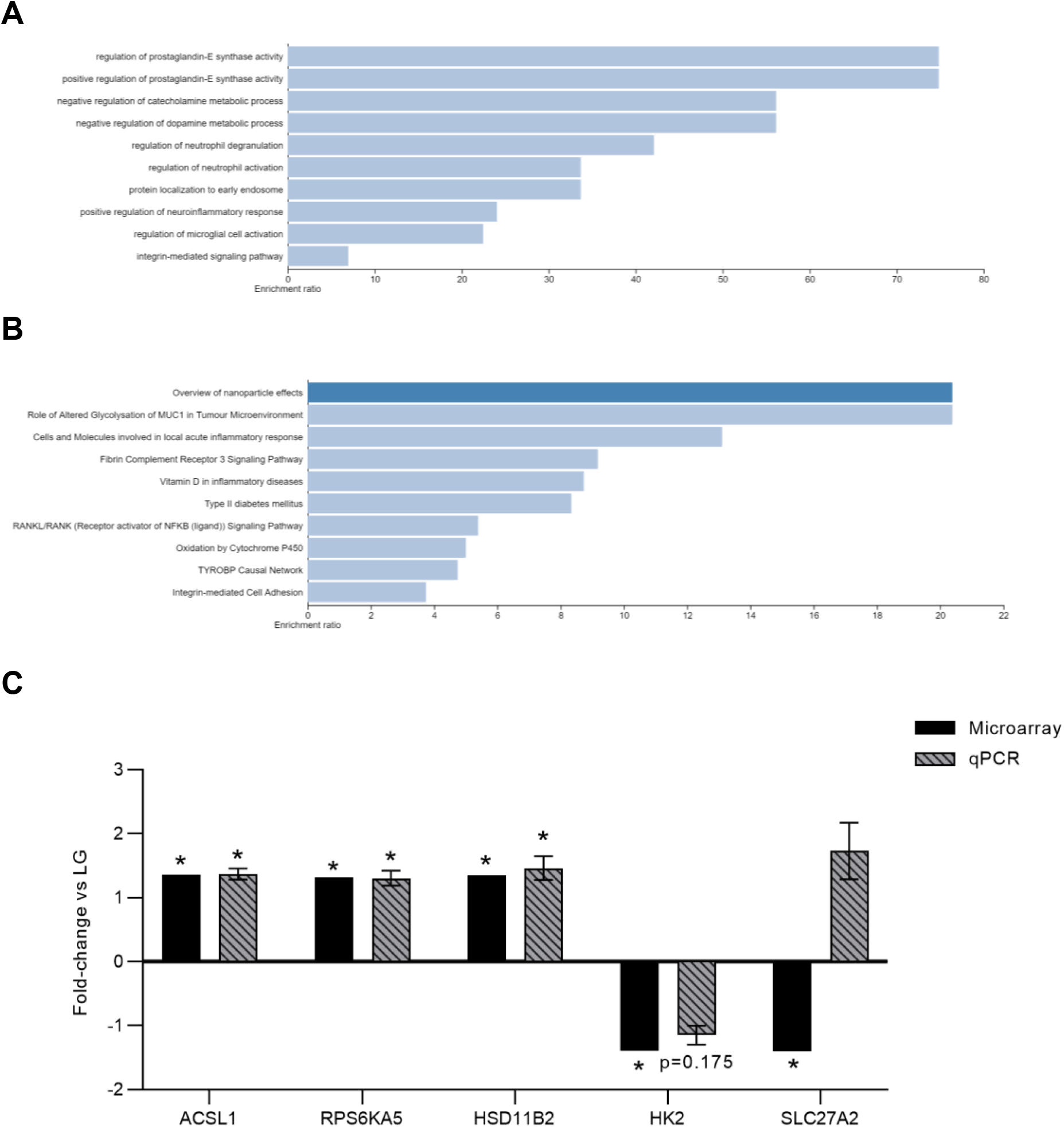
Pathway Analysis of Differentially expressed genes and RT-qPCR validation of microarray gene changes. Webgestalt was utilized to examine significantly enriched pathways of the differentially expressed genes identified in the Clariom S mRNA microarray. **(A)** Top 10 Geneontology biological pathways and **(B)** top 10 Wikipathways functional pathways enriched in the differentially expressed genes. Dark blue pathways were significantly enriched (FDR p < 0.05) and light blue pathways were not significantly enriched in the gene set (FDR p > 0.05). **(C)** RT-qPCR was utilized to validate differentially expressed genes identified in the microarray. Data was analyzed via paired t-test (n=8/group), and data expressed as fold-change vs LG BeWo CT (mean ± SEM). Down-regulated genes ratios were inversed and expressed as a negative (e.g 0.5-fold is expressed as a minus-2 fold-change).

As there were no enriched metabolism-related pathways, individual differentially expressed genes involved in cellular metabolic processes were individually identified and selected for validation via RT-qPCR. Specifically, ACSL1, RPS6KA5, HSD11B2, HK2 and SLC27A2 were selected for validation (Fig. 10C). The altered expression of ACSL1 (+1.36 FC in the microarray; +1.369 FC in the RT-qPCR validation); RPS6KA5 (+1.32 FC in the microarray; +1.304 FC in the RT-qPCR validation); and HSD11B2 (+1.35 FC in the microarray; +1.463 FC in the RT-qPCR validation) in HG-cultured BeWo CT cells were validated via RT-qPCR. (Fig. 10C) While not significant, the expression of HK2 was found slightly reduced in HG BeWo CT cultures via RT-qPCR (−1.152 FC vs LG BeWo CT cells). The differential expression of SLC27A2 in HG-cultured BeWo CT cells could not be validated via RT-qPCR (Fig. 10C).

### Impacts of HG culture conditions on the metabolome of BeWo CT cells

On average 6541 ± 42 (mean ± SD) metabolite peak pairs were measured in each sample. A summary of the metabolites identified in all tiers is available in **Supplementary Table 4.** Of these peak pairs, 179 were positively identified in tier 1 and 602 peak pairs were identified with high confidence in tier 2. Of these identified peak pairs, 7 from tier 1 and 116 from tier 2 were found to be differentially abundant (≥ ±1.5 FC, FDR-adjusted p<0.05) between HG and LG cultured BeWo CT cells. A list of all differentially abundant metabolites between LG and HG-cultured BeWo CT cells is available in **Supplementary Table 5**.

Differentially abundant metabolites were subsequently visualized via volcano plot (±1.5 FC, p<0.05) and heat map (Fig. 11A,B). Additionally, the degree of differences in metabolite profiles between HG and LG-cultured BeWo CT cells was visualized by unsupervised principal component analysis (PCA) 2D plot as well as supervised partial least squares discriminant analysis (PLS-DA) scores plot (Fig. 12A,B).

**Figure 11.**
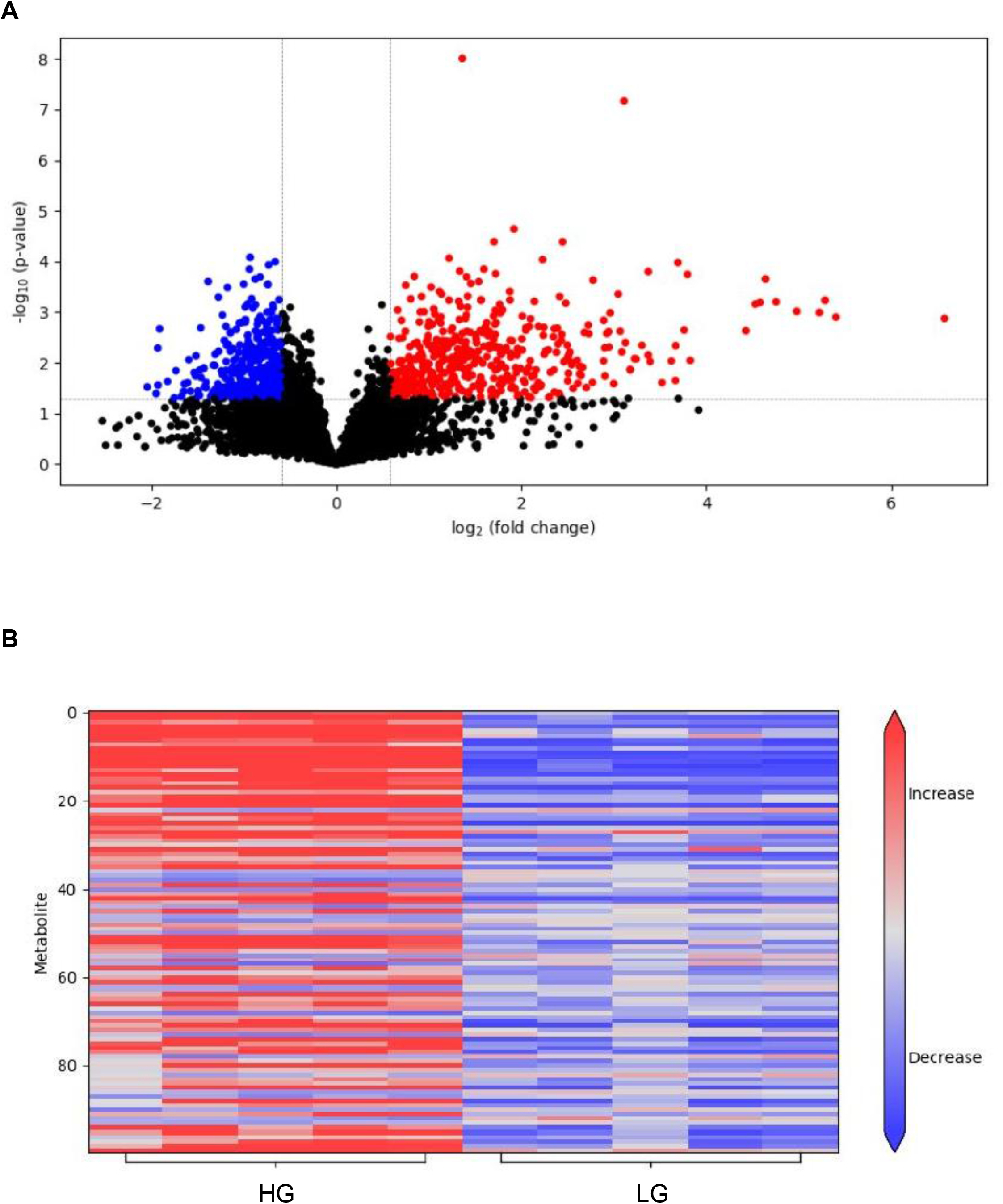
Visualization of differentially abundant metabolites in HG-cultured BeWo CT cells. **(A)** Volcano plot visualizing differentially abundant metabolites in HG-cultured BeWo CT cells ((≥ ±1.5-fold change, p<0.05, n=5/group). The x-axis indicated log_2_(fold-change) vs LG BeWo CTs, and the y-axis indicates the p-value (−log10). The red dots represent significantly increased metabolites, and the blue dots represent significantly decreased metabolites in HG-cultured BeWo CT cells. **(B)** heat map visualizing the top-100 (by p-value) differentially abundant metabolites in HG-cultured BeWo CT cells. Each column represents a different sample, and each row represents an individual differentially abundant metabolite. Metabolite abundance was color coded with red representing increased metabolite abundance and blue representing decreased metabolite abundance.

**Figure 12.**
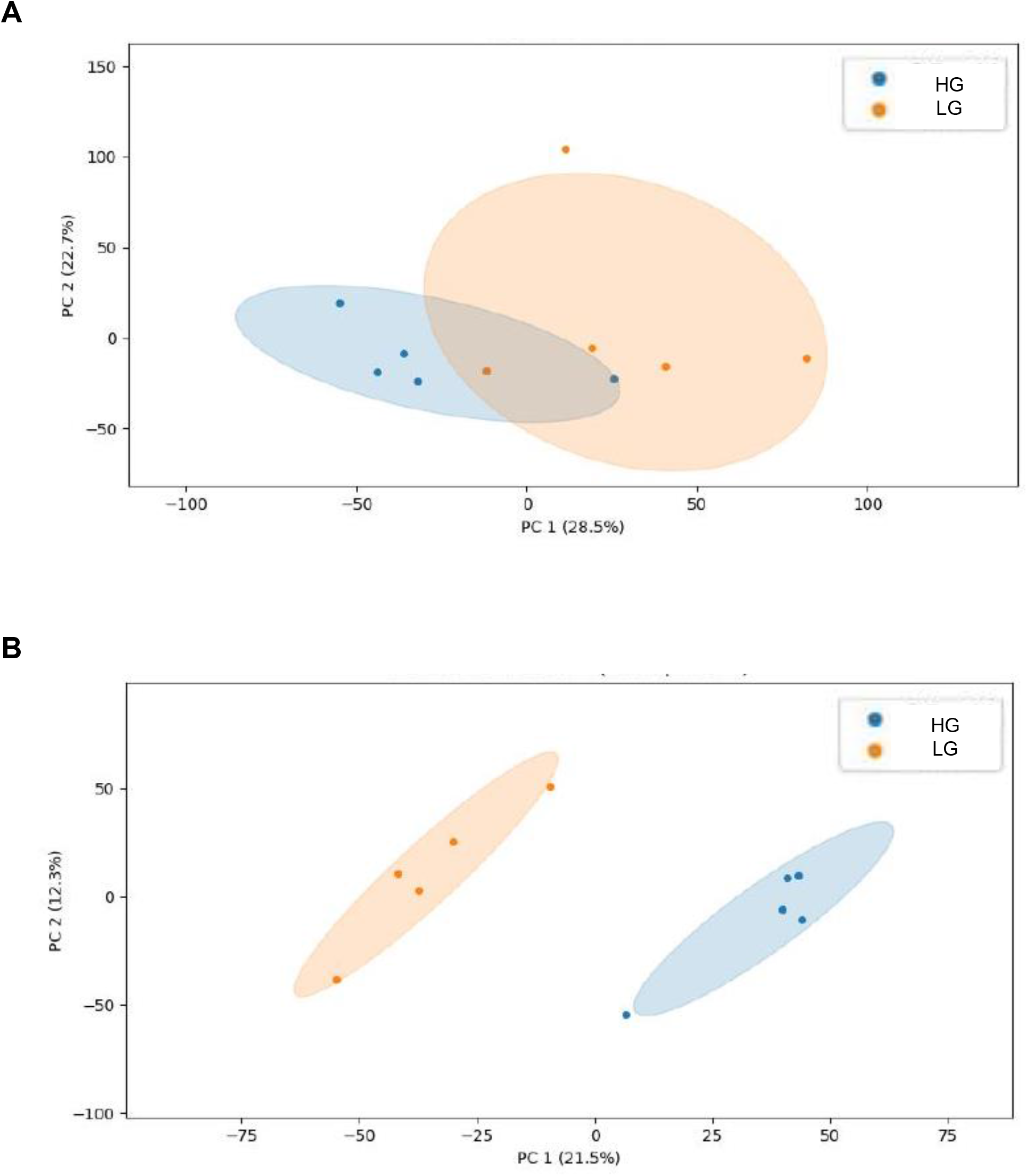
Visualization of the degree of separation between metabolite profiles in HG and LG cultured BeWo CT cells. (A) Unsupervised principal component analysis (PCA) and **(B)** supervised partial least squares discriminant analysis (PLS-DA) plots were constructed to visualize the separation in metabolite concentrations between LG and HG-cultured BeWo CT cells.

Pathway analysis revealed that the glycolysis/gluconeogenesis (significantly increased lactate (+2.72 FC) levels, and significantly decreased acetaldehyde (+2.72 FC) levels); pyruvate metabolism (increased lactate (+2.72 FC) levels, and decreased acetaldehyde (−1.20 FC) levels); drug metabolism – cytochrome P450 (increased 2-hydroxyiminostilbene (+2.52 FC) and 3-carbamoyl-2-phenylpropionic acid (+5.42 FC) levels); ascorbate and aldarate metabolism (increased L-gulonolactone (+2.46 FC) levels); riboflavin metabolism (increased riboflavin (+3.68 FC) levels); fatty acid biosynthesis (increased malonate (+3.74 FC) levels); as well as synthesis and degradation of ketone bodies (increased (R)-3-hydroxybutanoate (+1.60 FC) levels) pathways were significantly enriched in HG-cultured BeWo CT cells (FDR-corrected p<0.05). A scatterplot of the pathway nodes and summary of the differentially abundant metabolites within each pathway is available in Fig. 13.

**Figure 13.**
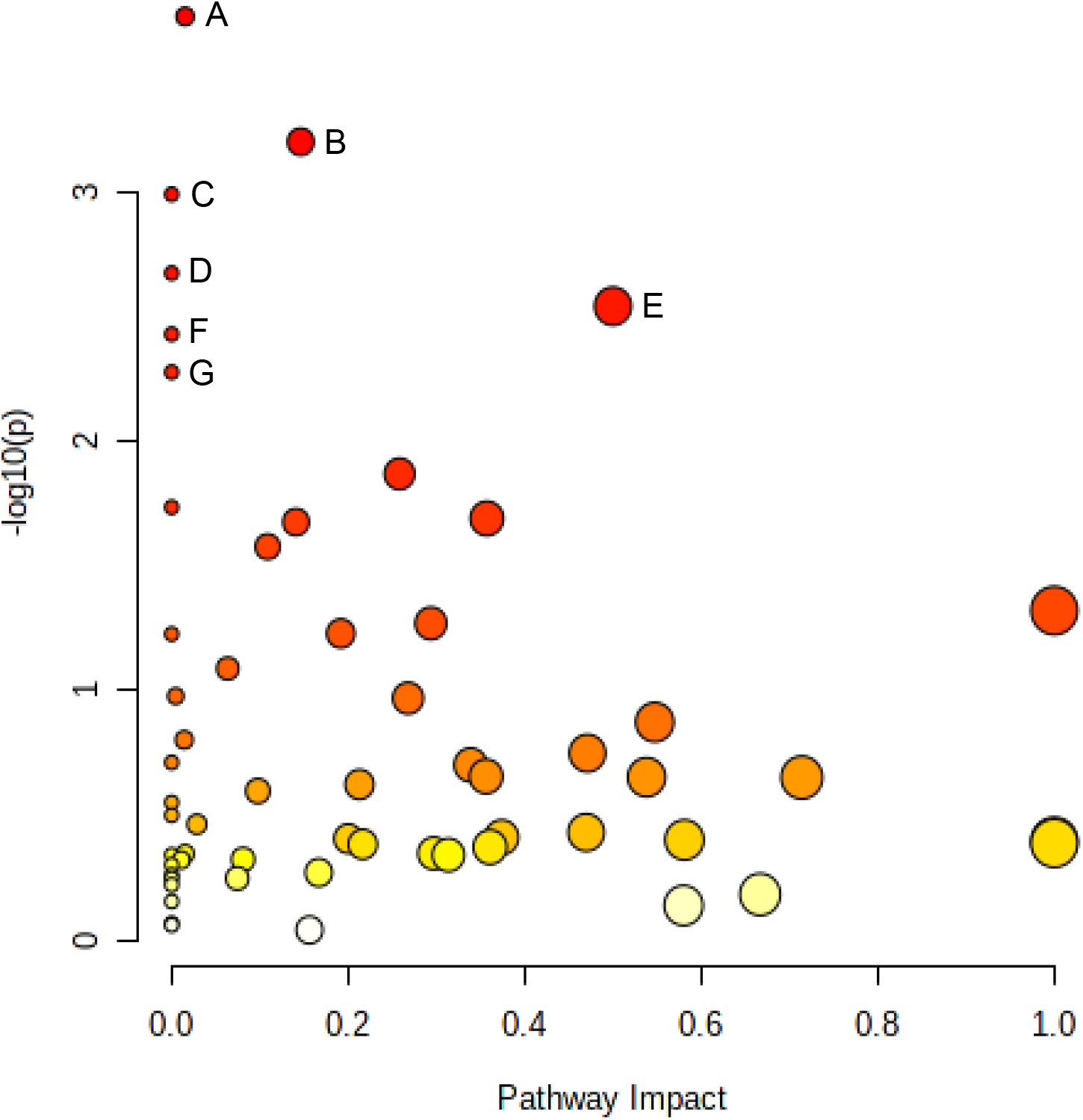
Pathway analysis of metabolites identified in tiers 1 and 2. Identified peak pairs from tiers 1 and 2 were imported into MetaboAnalyst v5.0 for analysis of enriched KEGG pathways. A scatterplot was created to visualize identified pathways with pathway impact on the x-axis and p-value (−log10) on the y-axis. Pathways with a false discovery rate p<0.05 were determined to be significant and were labelled with uppercase letters. The **(A)** Glycolysis/Gluconeogenesis, **(B)** Pyruvate Metabolism, **(C)** Drug Metabolism – Cytochrome P450, **(D)** Ascorbate and Aldarate Metabolism, **(E)** Riboflavin Metabolism, **(F)** Fatty Acid Biosynthesis, and **(G)** Synthesis and Degradation of Ketone Bodies pathways were significantly enriched in HG-cultured BeWo CT cells.

## Discussion

The current study aimed to expand upon current published literature [22,30,31,34] and more thoroughly explore the independent impacts of hyperglycemia on placental trophoblasts by characterizing nutrient storage and mitochondrial respiratory activity in BeWo trophoblasts following a relatively prolonged 72-hour HG-exposure (25 mM glucose). While previous studies utilizing BeWo trophoblasts have specifically highlighted the impacts of high glucose exposure on undifferentiated CT cells [30,31,34] the current study is strengthened through the combined examination of both undifferentiated CT cells and differentiated SCT cells as well as by the use of functional readouts of metabolic and mitochondrial activity. The more chronic 72-hour culture protocol as utilized in this study allowed for exposure of villous trophoblast cell populations to hyperglycemia prior to and during differentiation, analogous to *in vivo* villous trophoblast layer development whereby progenitor CT cells are pre-exposed to dietary nutrients prior to and during syncytialization. Subsequently, the current study utilized a multi-omics research approach and described altered transcriptomic and metabolic signatures in HG-exposed BeWo CT cells, that further demonstrated glucose-mediated alterations to placental metabolic function. Overall, the data presented in this study demonstrated that a 72-hour exposure to hyperglycemia, while sufficient to modulate transcriptomic and metabolomic signatures as well as increase triglyceride and glycogen nutrient stores in BeWo trophoblast cells, is not associated with direct impairments in mitochondrial respiratory activity.

### Hyperglycemia and nutrient stores in BeWo Trophoblasts

As previously highlighted, DM during pregnancy is associated with abberant nutrient storage in the villous trophoblast layer of the placenta [18–23]. Our results demonstrated that HG-culture conditions directly facilitate increased glycogen content in both BeWo CT and SCT cells, and additionally that CT cells have a greater glycogen storage potential than SCT cells. It is important to note that these differentiation state-dependent trends in glycogen content are consistent with previous reports from primary human placental tissue which described that glycogen storage predominately occurs within the CT cells of the placenta [24,46,47]. Increased glycogen abundance in diabetic placentae has been thought to be a mechanism by which the placenta limits materno-fetal glucose transfer in times of nutrient overabundance to limit fetal overgrowth [24, 25]. However, we speculate that the increased glycogen stores observed in the BeWo trophoblast cells also functions to limit the pool of glucose available for glycolysis and subsequent mitochondrial oxidation and in turn acts to protect trophoblast mitochondria from glucose-mediated oxidative damage.

High-glucose exposure in both BeWo CT and SCT cells was also associated with reduced relative protein abundance of glycogen synthase as well as with increased inhibitory Ser^641^ phosphorylation [48] of glycogen synthase. As there were no glucose-mediated differences in the protein abundance of GSK3β or the inhibitory Ser^9^ phosphorylation [49] of GSK3β, the increased inhibitory phosphorylation of glycogen synthase in HG-cultured BeWo trophoblasts likely occurs via a GSK3β-independent mechanism. These data could suggest that increased glycogen levels in placental trophoblasts act via a negative feedback mechanism to prevent excessive glycogen accumulation by altering glycogen synthase protein abundance and post-translational modifications, as has previously been described [50]. Overall, these data suggested that following 72-hours of hyperglycemia BeWo trophoblast cells have a diminished capacity to store excess glucose as glycogen. Thus, we speculate that HG-cultured BeWo trophoblast cells are at risk of developing glucose-toxicity if the exposure is continued beyond our 72-hour timepoint.

In addition to increased glycogen stores, our study also demonstrated increased triglyceride content in BeWo CT in response to excess glucose exposure. These data were consistent with previous reports that have highlighted an increased accumulation of triglyceride species in placental explants from healthy pregnancies under high glucose conditions [22]. The current study further identified that there were no differentiation state-dependent differences in triglyceride abundance in BeWo trophoblast cells, similar to what has been reported in freshly isolated villous trophoblast samples [51]. Readouts from primary placenta samples further highlighted that lipid esterification processes occur primarily within villous CT cells, and it was suggested that lipid droplets present in primary SCT cells could be remnants from esterification process that occurred prior to syncytialization [51]. We speculate that a similar reduction in esterification activity also occurs in BeWo SCT cells and may explain the absence of a HG-mediated difference in triglyceride content in our BeWo SCT cells. Future studies utilizing functional readouts of lipid esterification activity may be needed be better elucidate the mechanisms underlying differences in TG abundance between undifferentiated BeWo CT cells and differentiated BeWo SCT cells.

Overall, these data suggest that hyperglycemia is an independent modulator of nutrient storage in trophoblast cells and highlights that increased glucose availability may be a direct mechanism underling abberant placental nutrient storage in diabetic pregnancies.

### Hyperglycemia and metabolic function in BeWo Trophoblasts

The current study demonstrated that cellular mitochondrial respiratory activity (measured by the Seahorse XF Mito Stress Test) as well as that the activities of ETC complexes I and II are not impacted in BeWo trophoblasts cultured under hyperglycemia at 72H. Additional analysis of metabolic function via the Seahorse Glycolysis Stress Test and activity assays for LDH and CS enzymes further indicated that isolated hyperglycemia does not impact functional aspects of metabolism in BeWo trophoblasts. Overall, our data suggests that hyperglycemia alone may not directly facilitate the impairments in mitochondrial respiratory activity that have previously been observed in DM-exposed primary placental samples [27–29].

Previous studies in other cell preparations, however, have demonstrated that the impacts of HG-culture conditions on mitochondrial respiratory function are dependent on the length of high-glucose exposure. For example, human kidney tubule (HK-2) cells displayed reduced basal and maximal mitochondrial respiratory activity only when cultured under HG-conditions (25 mM) for at least 4 days [52]. In contrast, mitochondrial respiratory activity in kidney glomerular (HMC) cells was not impacted prior to 8 days of high glucose exposure [52]. Likewise, mitochondrial activity of human umbilical cord endothelial (EA.hy926) cells was not impaired after 3 days of high glucose (25 mM) treatment, but was reduced after 6 days of HG-culture conditions, and this impairment in mitochondrial function was sustained through 9 days of high glucose exposure [53]. Overall, these studies suggest that there are time-course dependent factors that influence whether hyperglycemia impacts cellular mitochondrial respiratory activity. Thus, the conclusions of the current study may be limited due to the single timepoint utilized for all analyses of metabolic function. It is possible that prolonging HG-culture conditions in BeWo trophoblasts could ultimately lead to impaired mitochondrial function, as was observed in EA.hy926 and HMC cells, and the impacts of a longer duration of high glucose treatment in BeWo cells may need to be explored in future investigations.

In addition to impairing mitochondrial respiratory activity in trophoblast cells, previous research has highlighted that hyperglycemia negatively regulates mitochondrial function in some cell types through altering mitochondrial fusion (regulated by OPA1) and fission (regulated by DRP1) dynamics leading increased mitochondrial fractionation [54–57]. Increased mitochondrial fission is associated with increased cellular oxidative stress, and impaired insulin sensitivity, and ultimately mitochondrial respiratory dysfunction [58, 59]. We observed a trend towards increased (mean 19% increase) pSER^616^ phosphorylation of DRP1 (a post-translational modification associated with increased mitochondrial translocation of DRP1 and subsequently increased mitochondrial fission) in HG-cultured BeWo trophoblasts [60]. These post-translational modifications of DRP1, although non-significant, may indicate that important underlying aspects of trophoblast mitochondrial function are negatively regulated by hyperglycemia, even though global readouts of BeWo mitochondrial respiratory activity were not impacted. We speculate that this could indicate that the HG-cultured BeWo trophoblasts are at an early timepoint in a transition towards mitochondrial dysfunction.

### Transcriptomic analysis of HG-cultured BeWo CT cells

While functional readouts of BeWo trophoblast mitochondrial function were sustained in response to hyperglycemia, we did observe impaired mitochondrial respiratory activity in differentiated BeWo SCT cells. Syncytialization of BeWo trophoblasts was associated with reduced mitochondrial spare respiratory capacity, reduced coupling efficiency, concomitant with reduced activity of ETC complex II and reduced protein expression of ETC complex IV. Furthermore, BeWo SCT cells displayed increased DRP1 protein abundance and decreased OPA1 protein abundance suggestive of increased mitochondrial fractionation. These data may indicate that alterations in mitochondrial dynamics underlies the observed functional differences in mitochondrial respiration between BeWo CT and SCT cells. Previous work by our research group has likewise highlighted that BeWo CT cells are overall more metabolically active than BeWo SCT cells [37], and similar trends have been reported in cultured PHT cells [28, 61]. As undifferentiated BeWo CT cells display greater metabolic activity and greater alterations to nutrient stores in response to hyperglycemia than in differentiated SCT cells, we speculated that alterations in transcriptome and metabolome profiles in response to high glucose culture conditions would be more prevalent in these progenitor cells. Thus, this current study sought to examine global gene expression as well as global metabolite abundance solely in HG-cultured BeWo CT cells to further elucidate mechanisms underlying altered placental metabolic function in response to hyperglycemia.

The current study identified 197 differentially expressed genes (≥ ±1.3 FC) in BeWo CT cells cultured under hyperglycemia for 72H. However, previous reports in BeWo CT cells demonstrated more substantial variations in gene expression between BeWo trophoblasts cultured under similar high and low glucose conditions [30]. The differences between the current study and previous reports may be due in part to differences in study design (pooling samples for arrays vs independent arrays for each sample), cell media formulation (DMEM-F12 vs F12K) as well as length of hyperglycemic exposure (48H vs 72H). Despite these differences in the number of differentially expressed genes, our study did align with this previous transcriptomic report, and further demonstrated that elevated glucose levels impact the expression of genes involved in metabolic processes [30]. Of note, HG-culture conditions in the current study were associated with increased mRNA expression of *ASCL1* in BeWo CT cells. Additionally, we observed a simultaneous trend towards increased protein abundance of *ACSL1*, although these trends did not reach statistical significance. Previous studies have highlighted that *ACSL1* is involved in lipid synthesis in various tissues and that knockdown of *ACSL1* is associated with reduced triglyceride and lipid droplet abundance [62–65]. More importantly, cells transfected to overexpress *ACSL1* have been found to have increased triglyceride accumulation [65–67]. Thus, we speculate that *ACSL1* may be involved in the trafficking of lipid species to lipid droplets and may underlie the glucose-induced accumulation of triglyceride species that was observed in our BeWo CT cultures.

Our data additionally highlighted that isolated hyperglycemia in BeWo CT cells was associated with an increased expression of *HSD11B2*, an enzyme involved in the metabolism and inactivation of the glucocorticoid cortisol. In normal pregnancies, cortisol is thought to be involved in mediating the physiological increase in maternal insulin resistance that is necessary to support fetal growth [68, 69], however, circulating cortisol levels may be pathologically elevated in some GDM pregnancies [70]. Interestingly, placentae from GDM pregnancies have been found to have increased expression of *HSD11B2* leading to increased cortisol inactivation [71]. As, elevated cortisol levels in fetal circulation have been associated with impaired brain development processes [72], increased placental *HSD11B2* expression in GDM pregnancies may act in a protective manner to limit fetal glucocorticoid exposures. However, as cortisol also activates the glucocorticoid receptor leading to modulation of gene transcription [73, 74], alterations in placental cortisol metabolism may have downstream consequences on placental function through altering gene expression. Overall, the results from the current study demonstrated that hyperglycemia is an important regulator of placental *HSD11B2* expression. However, it remains poorly understood whether these glucose-mediated impacts to placental cortisol metabolism act in a protective or detrimental manner.

### Metabolomic analysis of HG-cultured BeWo CT cells

Multivariate analysis of BeWo trophoblast metabolome profiles highlighted a divergence in metabolite signatures between HG and LG-cultured BeWo CT cells suggesting that high glucose levels also impact metabolite levels in the placenta. Subsequent pathway analysis highlighted specific intracellular accumulations of lactate (involved in the glycolysis/gluconeogenesis and pyruvate metabolism pathways), malonate (involved in the fatty acid biosynthesis pathway), as well as riboflavin (involved in the riboflavin metabolism pathway) in the HG-cultured BeWo CT cells.

The observed lactate accumulation likely suggests that glycolytic flux is in fact increased in HG-cultured BeWo CT cells [75]. It is interesting to note that this study did not observe increased basal or maximal glycolytic activity when assessed via the Seahorse XF Glycolysis Stress Test. As the Glycolysis Stress Test utilizes extracellular media acidification (resulting from the co-export of lactate and H^+^ from the cell) as a proxy measurement of glycolytic activity, this functional assay may underestimate glycolytic activity in the event of reduced lactate export as could potentially occur in a “Cytosol-to-Mitochondrial Lactate Shuttle” metabolic pathway [75].

Increased malonate levels may reflect an increase in *de novo* lipogenesis via FASN in HG-cultured BeWo CT and may be another mechanism underlying the increased TG levels observed in HG-cultured BeWo CT cells [76]. Future studies and the use of radio-labelled metabolites may be required to further characterize glycolytic activity and *de novo* lipogenesis in high-glucose exposed BeWo trophoblasts.

The accumulation of riboflavin (the essential vitamin B_2_) in HG-cultured BeWo CT cells either reflects an increased cellular uptake of riboflavin or an inhibition of riboflavin metabolism into the cofactors flavin adenine dinucleotide (FAD) and flavin mononucleotide (FMN) [77]. Reduced metabolism of riboflavin to its cofactor intermediaries has previously been implicated in the development of mitochondrial dysfunction [78]. However, riboflavin has also previously been suggested to act as an antioxidant [79] and has been demonstrated to be beneficial in reducing oxidative stress in rodent models of DM [80, 81]. Future investigations may be needed to assess the impacts of riboflavin accumulation in HG-cultured trophoblasts and elucidate whether accumulation of this vitamin is beneficial or harmful to the placenta.

## Conclusion

The results of the current study highlighted that a 72-hour hyperglycemia exposure independently impacts metabolic function and nutrient storage in BeWo trophoblasts but does not mediate any global changes in functional readouts of mitochondrial respiratory activity. Interestingly, HG-cultured BeWo trophoblasts displayed markers of reduced glycogen storage capacity that potentially increases the supply of free glucose for oxidation, as well as markers suggestive of a transition towards altered mitochondrial dynamics, that overall may be indicative of early metabolic adaptations, or of more concern, a transition towards mitochondrial failure. It is important to note that while the 72-hour hyperglycemic exposure utilized in the current study is relatively prolonged in the setting of *in vitro* cell culture experiments, this timeline is acute in comparison to the 40-week duration of human pregnancy *in vivo*. Thus, preventing even short periods of hyperglycemia may be important in the clinical management of diabetic pregnancies to ensure appropriate placental function is sustained throughout gestation, and in turn that the risk of the offspring developing later life non-communicable metabolic diseases is limited.

## Data Availability Statement

Microarray data is available on the National Center for Biotechnology Information (NCBI) Gene Expression Omnibus (GEO) database (GSE190025). A full list of the metabolome peak pairs, and their high confidence and putative identifications are included in **Supplementary Table 4**.

## Author Contribution Statement

ZE and TR designed the study and drafted the manuscript. ZE was responsible for performing the experiments and all statistical analysis. XL and LL contributed to the metabolomics analysis. All authors read and approved the final manuscript.

## Acknowledgements

The authors would like to thank Dr. Ousseynou Sarr for his guidance and assistance in analyzing the microarray data presented in this manuscript.

