## Supplementary Tables 1-2; Supplementary Figures 1-9 for "The impact of hyperglycemia upon BeWo trophoblast cell metabolic function: A multi-OMICS and functional metabolic analysis"

**Supplementary Table 1.** Forward and reverse primer sequences used for quantitative Real-Time PCR analysis of BeWo cell syncytialization and mRNA microarray validation

| Gene | Accession No. | Annealing Temperature (°C) | Forward Sequence (5'-3') | Reverse Sequence (5'-3') | Efficiency | Amplicon Length (bp) |
|---|---|---|---|---|---|---|
| ACTB | NM_001101.4 | 60 | GTTGCTATCCAGGCTGTGCT | AGGTAGTCAGTCAGGTCCCG | 92.6% | 167 |
| PSMB6 | NM_002798 | 60 | CGGGAAGACCTGATGGCGGGA | TCCCGGAGCCTCCAATGGCAAA | 108.4% | 124 |
| CGB | Malhotra *et al* (2015) | 60 | CCCCTTGACCTGTGATGACC | TATTGTGGGAGGATCGGGGT | 105.3% | 120 |
| OVOL1 | Kusama *et al* (2018) | 60 | AGACATGGGCCACTTGACAG | AGGTGAACAGGTCTCCACTG | 101.1% | 104 |
| SLC27A2 | NM_003645.4 | 60 | ACTTCTGGAACCACAGGTCTTC | TCATCTGCCTTCAATCCGCTT | 107.2% | 101 |
| HSD11B2 | NM_000196.4 | 60 | GGCTGCTTCAAGACAGAGTCAG | GCTCGATGTAGTCCTTGCCG | 100.9% | 118 |
| ACSL1 | NM_001995.5 | 60 | AGTCAATCCTTGCCCAGATGA | ATCCGGTTCAGCAGTCTTGG | 103.8% | 198 |
| RPS6KA5 | NM_001322232.2 | 60 | GTGCCTGCACCATTTAAGCC | AGCAACAAAGGAATAGCCCTGA | 95.0% | 147 |
| HK2 | NM_000189.5 | 60 | ACGCCAAAATCACGTCTCCG | AGAGATACTGGTCAACCTTCTGC | 98.8% | 283 |

**Supplementary Table 2:** Specifications of antibodies utilized in immunoblotting experiments, and the protein mass loaded for each protein target

| Protein Target | Source | Dilution | Blocking solution | Protein Loaded | Heat (°C) | Company | Catalogue No. |
|---|---|---|---|---|---|---|---|
| Human Mitoprofile | Mouse Monoclonal | 1:1000 | 5% Milk | 35 µg | 37 | ABCAM | ab110411 |
| Glycogen Synthase | Rabbit Polyclonal | 1:1000 | 5% BSA | 25 µg | 95 | Cell Signaling Technologies | 3893 |
| $pSer^{641}$ Glycogen Synthase | Rabbit Polyclonal | 1:1000 | 5% BSA | 20 µg | 95 | Cell Signaling Technologies | 3891 |
| GSK3-β | Rabbit Monoclonal | 1:1000 | 5% BSA | 25 µg | 95 | Cell Signaling Technologies | 9315 |
| pSer9 GSK3-β | Rabbit Monoclonal | 1:1000 | 5% BSA | 20 µg | 95 | Cell Signaling Technologies | 9323 |
| ACSL1 | Rabbit Polyclonal | 1:500 | 5% BSA | 12.5 µg | 95 | Cell Signaling Technologies | 4047 |
| Fatty Acid Synthase | Rabbit Monoclonal | 1:1000 | 5% BSA | 15 µg | 95 | Cell Signaling Technologies | 3180 |
| OPA1 | Rabbit Monoclonal | 1:1000 | 5% BSA | 15 µg | 95 | Cell Signaling Technologies | 67589 |
| DRP1 | Rabbit Monoclonal | 1:1000 | 5% BSA | 15 µg | 95 | Cell Signaling Technologies | 8570 |
| pSER616 DRP1 | Rabbit Monoclonal | 1:1000 | 5% BSA | 12.5 µg | 95 | Cell Signaling Technologies | 4494 |
| GLUT1 | Mouse Monoclonal | 1:1000 | 5% BSA | 15 µg | 20 | EMD Millipore | mabs132 |
| Anti-Rabbit Secondary | Goat | 1:10,000 | - | - | - | Cell Signaling Technologies | 7074 |
| Anti-Mouse Secondary | Horse | 1:10,000 | - | - | - | Cell Signaling Technologies | 7076 |

A

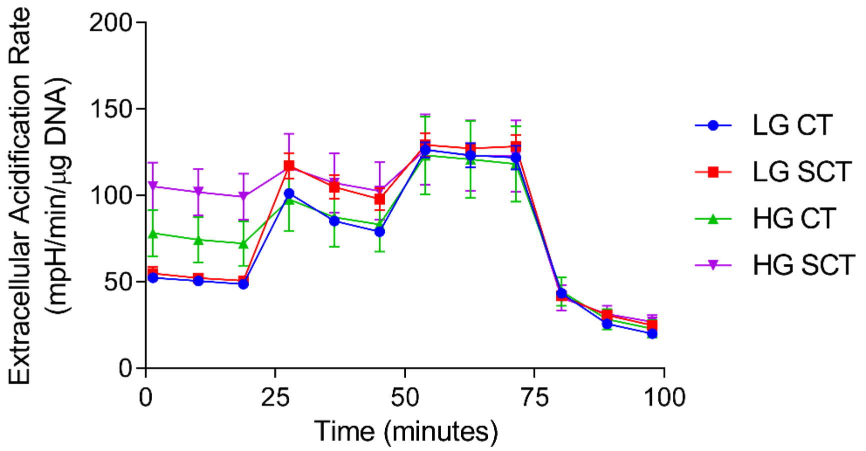


B

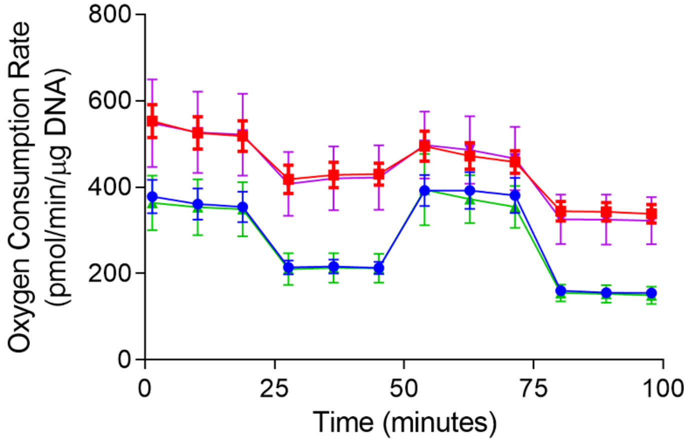


**Supplementary Figure 1 (A)** Representative Seahorse XF Glycolysis Stress Test assay tracings; data is presented as mean Extracellular Acidification Rate (ECAR) ± SEM of the technical replicates of each treatment group from one experiment. **(B)** Representative Seahorse XF Mito Stress Test assay tracings; data is presented as mean Oxygen Consumption Rate (OCR) ± SEM of the technical replicates of each treatment group from one experiment.

A

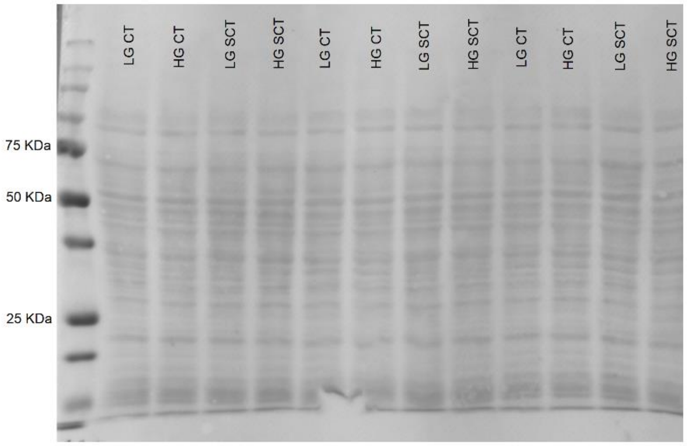


B

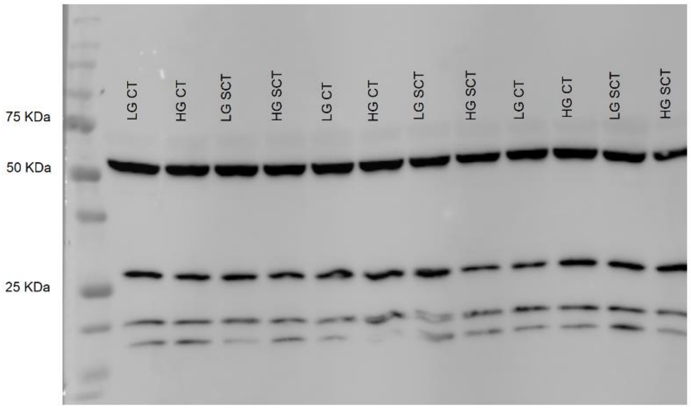


C

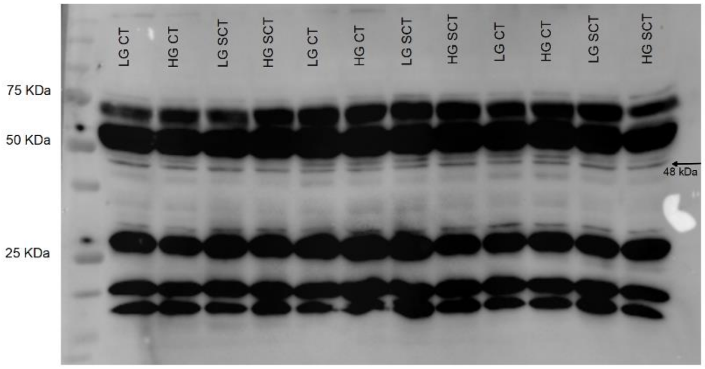


**Supplementary Figure 2. Full length representative images of Mito Profile immunoblots. (A)** representative full-length ponceau image used to quantify total lane protein; **(B)** representative full length blot used to quantify band abundance for ETC complexes I (18 kDa), II (29 kDa), IV (22 kDa), and V (54 kDa) (9-second ChemiDoc exposure); **(C)** representative full length blot used to quantify band abundance for ETC complex III (48 kDa) (140-second ChemiDoc exposure).

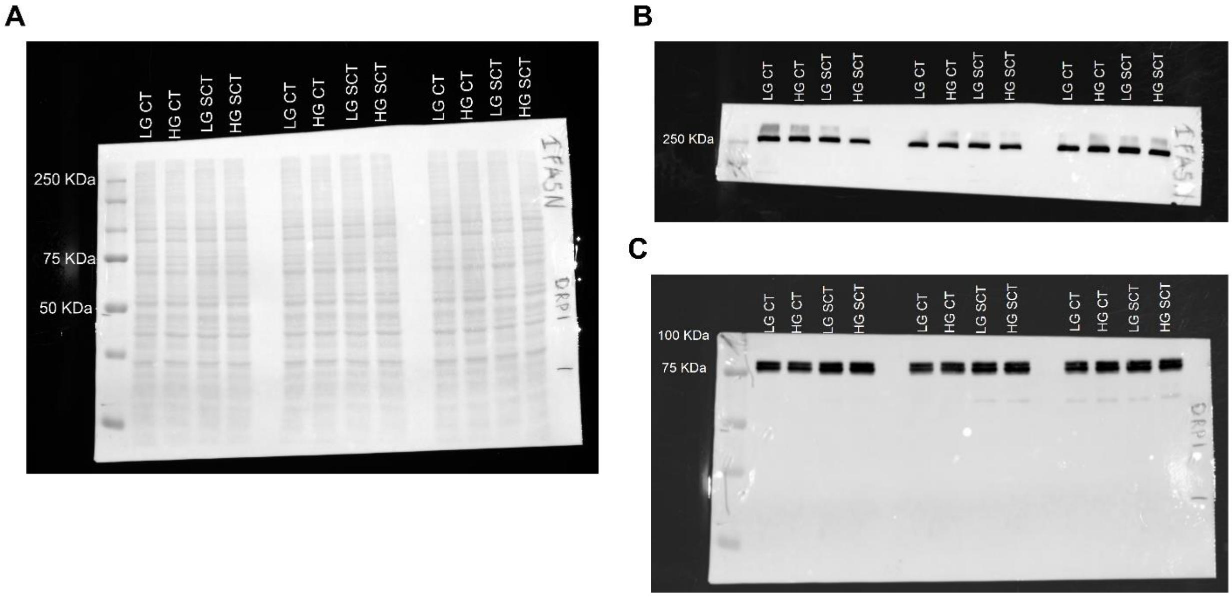


**Supplementary Figure 3. Uncropped representative images of total DRP1 and FASN immunoblots.** **(A)** Representative full length ponceau used to quantify total lane protein; **(B)** uncropped FASN bands; **(C)** uncropped total DRP1 bands.

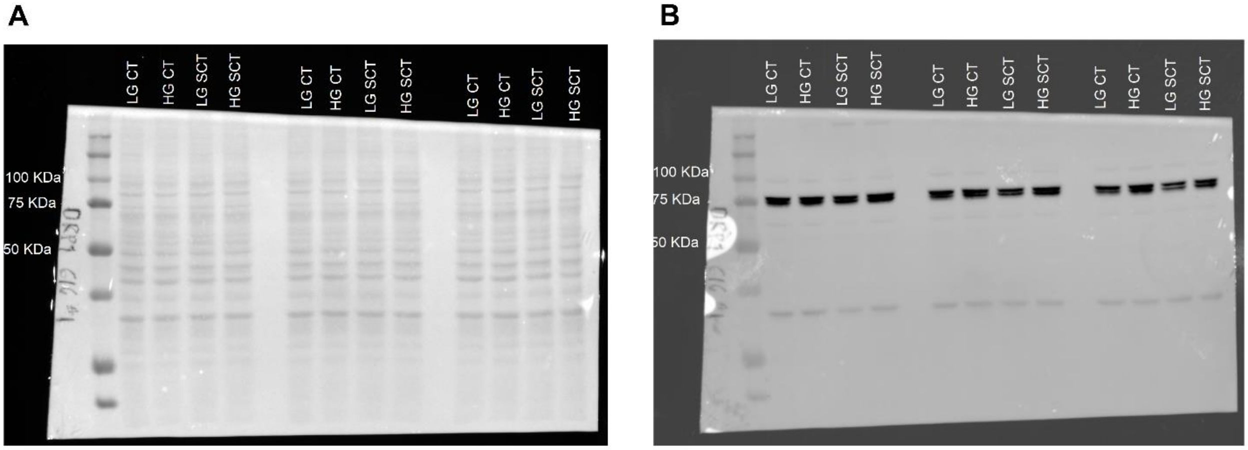


**Supplementary Figure 4. Uncropped representative images of pSER$^{616}$ DRP1 immunoblots. (A)** Representative full length ponceau used to quantify total lane protein; **(B)** uncropped pSER$^{616}$ DRP1 bands.

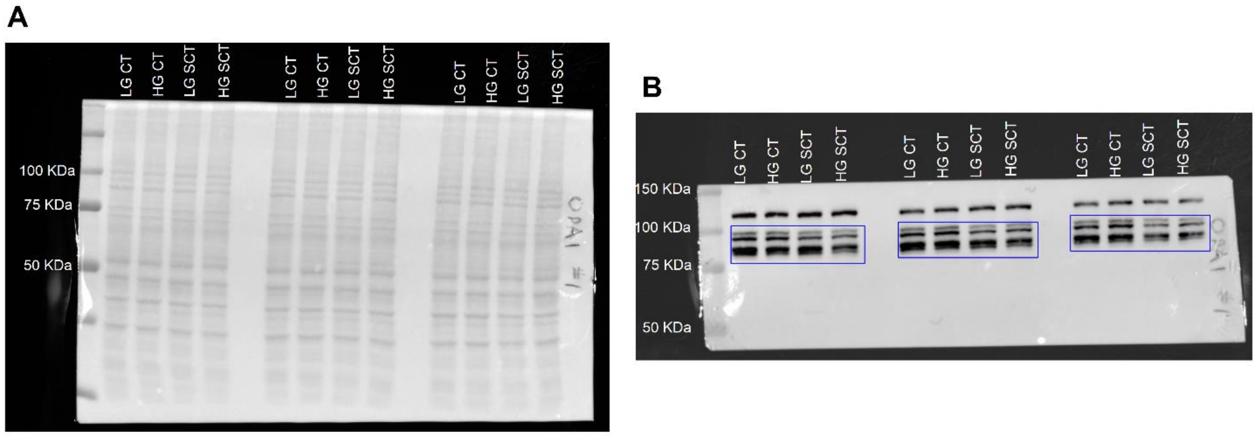


**Supplementary Figure 5. Uncropped representative images of OPA1 immunoblots. (A)** Representative full length ponceau used to quantify total lane protein; **(B)** uncropped OPA1 bands. Bands between 80-100 KDa were utilized to quantify OPA1 protein abundance and are highlighted by blue boxes

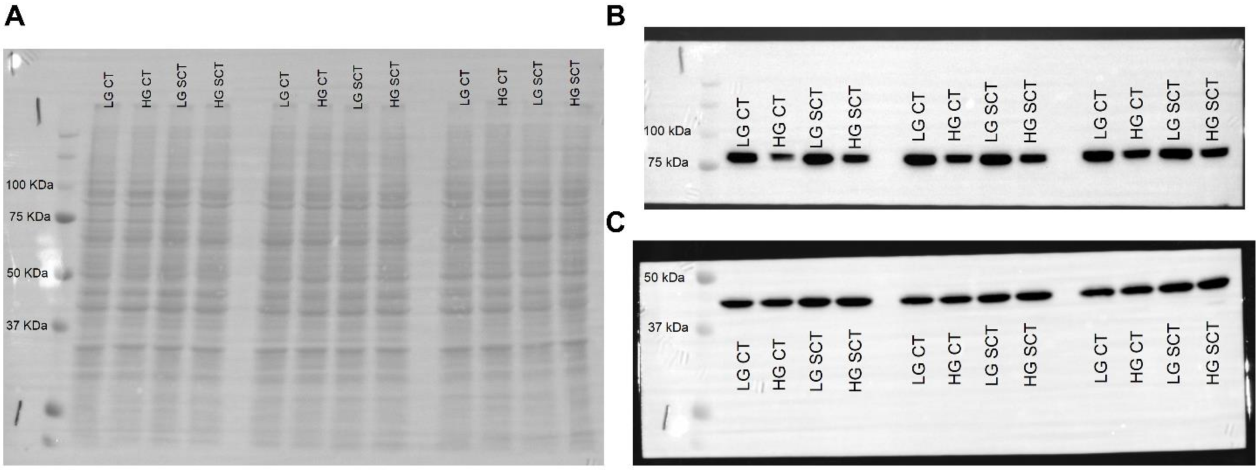


**Supplementary Figure 6. Uncropped representative images of Total Glycogen Synthase and GSK3β immunoblots. (A)** Representative full length ponceau used to quantify total lane protein; **(B)** uncropped total Glycogen synthase bands; **(C)** uncropped total GSK3β bands.

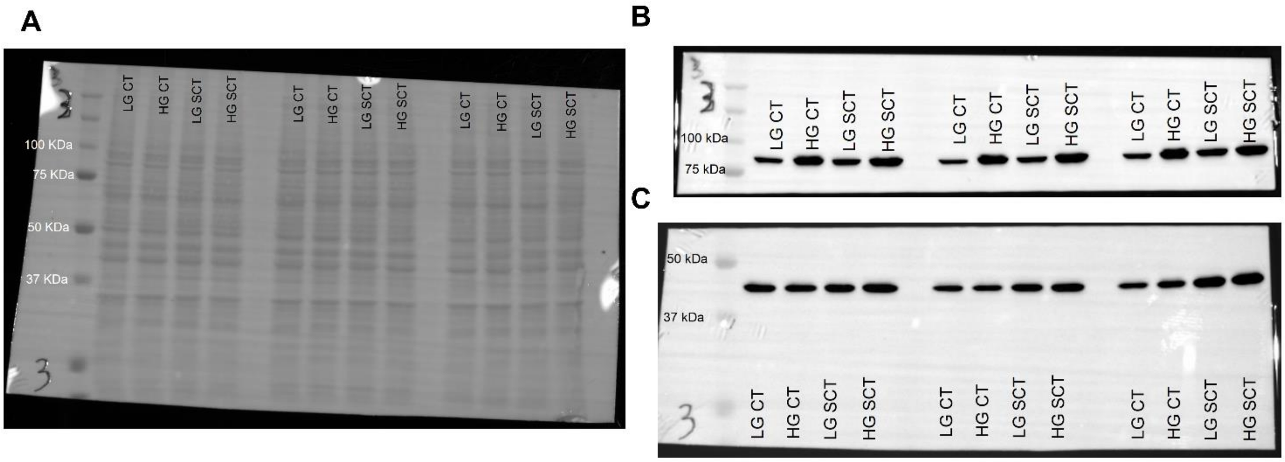


**Supplementary Figure 7. Uncropped representative images of pSer$^{641}$ Glycogen Synthase and pSer$^{9}$ GSK3β immunoblots. (A)** Representative full length ponceau used to quantify total lane protein; **(B)** uncropped pSer$^{641}$ Glycogen synthase bands; **(C)** uncropped pSer$^{9}$ GSK3β bands.

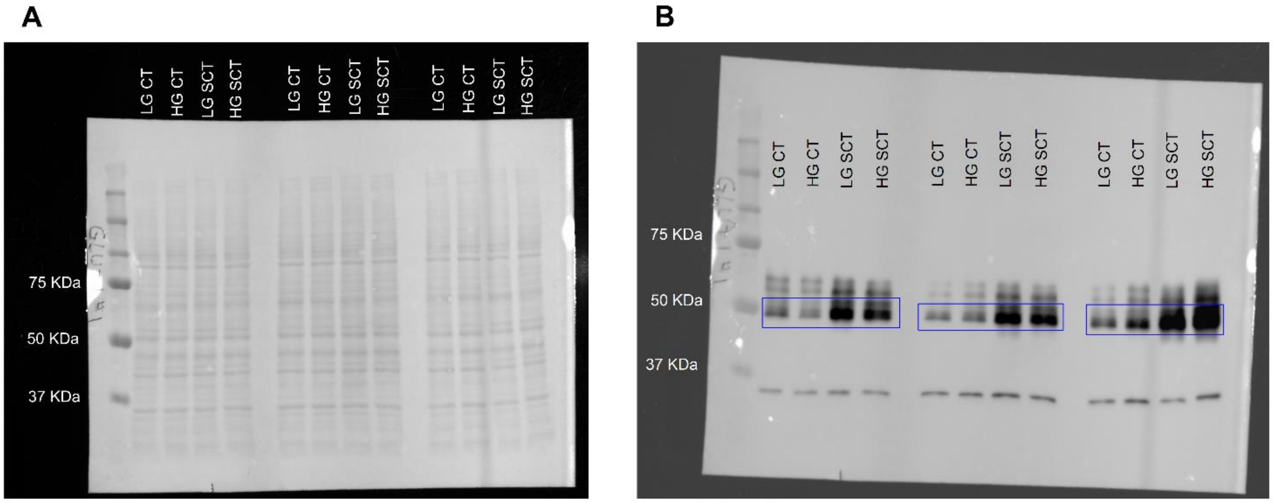


**Supplementary Figure 8. Uncropped representative images of GLUT1 immunoblots. (A)** Representative full length ponceau used to quantify total lane protein; **(B)** uncropped GLUT1 bands. Bands between 45-55 KDa were utilized to quantify GLUT1 protein abundance and are highlighted by blue boxes

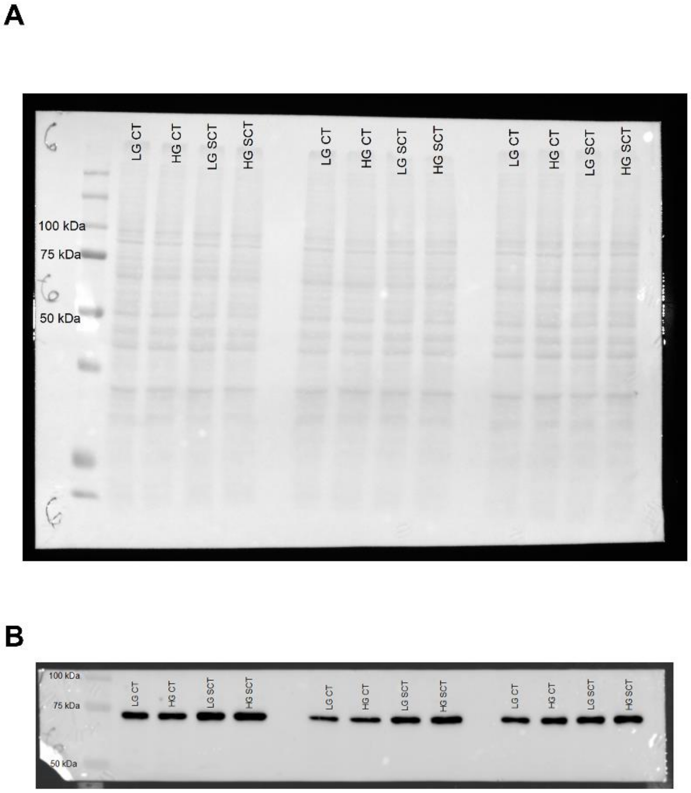


**Supplementary Figure 9. Uncropped representative images of ACSL1 immunoblots.** (A) Representative full length ponceau used to quantify total lane protein; (B) uncropped ACSL1 bands.
